# Vesicular pseudopodia define the fusion site on large secretory vesicles of the *Drosophila* salivary glands

**DOI:** 10.64898/2026.06.02.729163

**Authors:** Nadav Scher, Tom Biton, Vishnu Mohan, Neta Varsano, Noa Aharoni, Shari Carmon, Kamalesh Kumari, Eyal Schejter, Tamar Geiger, Yael Elbaz-Alon, Ori Avinoam

## Abstract

Large secretory vesicles (LSVs) pose a scaling problem for regulated exocytosis. Their micron-scale dimensions greatly increase the vesicular membrane surface area, making productive engagement between the vesicular and target membrane fusion machinery unlikely. Here, we show that vesicular pseudopodia define the fusion sites of LSVs in *Drosophila* larval salivary glands. Focused ion beam scanning electron microscopy revealed that most LSVs project polarized pseudopodia that interconnect neighboring vesicles and orient toward the apical membrane. Exposed pseudopodia were frequently observed at the apical surface and associated with narrow fusion pores, indicating that fusion occurs at these structures. Three-dimensional correlative light and electron microscopy showed that the I-BAR protein Missing in Metastasis (MIM) selectively localizes to exposed pseudopodia. Proteomic analysis based on a MIM pull-down assay identified exocyst components, including Sec15, which localizes to pseudopodia and persists at fusion sites throughout secretion. Finally, the tetraspanin Tsp42Ee marked complementary apical fusion domains and was required for efficient exocytosis. Our findings support a model in which prepatterned vesicular and apical membrane domains coordinate efficient exocytosis.

**Summary:** Regulated exocytosis of large secretory vesicles is facilitated by vesicular pseudopodia and an apical fusion domain that spatially organizes membrane tethering and fusion during secretion.

## Introduction

Regulated exocytosis is the process by which membrane-bound secretory vesicles release their luminal cargo after fusing with the plasma membrane. Fusion is mediated by conserved membrane-anchored SNARE proteins and their regulators, which bring vesicular and target membranes into close apposition and drive bilayer fusion (Jahn and Scheller, 2006; Söllner et al., 1993; Südhof and Rothman, 2009; Weber et al., 1998). Although the core fusion machinery is broadly conserved, the physical context in which it operates varies widely between secretory cells. In particular, many exocrine and other specialized secretory cells rely on large secretory vesicles (LSVs) with diameters in the micrometer range, rather than on the sub-micron vesicles that mediate neuronal and endocrine secretion (Jerdeva et al., 2005; Kliewer et al., 1985; Nightingale et al., 2012; Rousso et al., 2016; Wei et al., 2023).

Micrometer-scale vesicles impose distinct geometric and mechanical constraints on secretion. Their membrane area is substantially larger, their cargo is often dense or viscous, and secretion must occur while preserving the architecture and homeostasis of the apical surface. Consistent with this, studies in mammalian exocrine tissues and in the *Drosophila* larval salivary gland have shown that exocytosis of LSVs necessitates the recruitment of a specialized actomyosin meshwork that remodels the fused vesicle membrane and promotes efficient cargo release (Ebrahim et al., 2019; Kamalesh et al., 2021; Kamalesh et al., 2024; Masedunskas et al., 2011; Miklavc et al., 2015; Rousso et al., 2016; Tran et al., 2015).

A long-standing but unresolved idea is that part of this scaling problem is solved by pre-patterning a dedicated fusion domain on the vesicular membrane. Classical ultrastructural studies described protrusive structures extending from secretory granules, termed vesicular pseudopodia, and proposed that fusion with the cell surface may occur at these structures (Castel and Selinger, 1977; Schramm et al., 1972). These early studies also suggested that pseudopodia are regulated features of secretory granules, as their appearance increased under secretion-inducing conditions and they appeared oriented towards the apical surface (Schramm et al., 1972; Selinger et al., 1974). More recent synthesis of this literature has revived the possibility that vesicular pseudopodia are a conserved adaptation of large secretory granules (Scher and Avinoam, 2025). However, their mechanistic significance remains unknown. In particular, it remains unclear how prevalent pseudopodia are and whether they constitute *bona fide* pre-fusion membrane domains that spatially define the fusion site.

The *Drosophila* larval salivary gland provides a powerful system for addressing this question. During late larval development, adhesive mucin-like (“glue”) proteins are packaged into LSVs that undergo stereotyped biogenesis and maturation before an ecdysone-triggered burst of apical secretion (Biyasheva et al., 2001; Burgess et al., 2011; Ma et al., 2020; Neuman et al., 2021). This system has yielded mechanistic insight into vesicle formation, endosomal contribution to maturation, Rab-dependent identity and exocytosis, and exocyst-dependent regulation of both maturation and secretion (Freire et al., 2024; Ma et al., 2020; Neuman et al., 2021). Importantly, mature glue LSVs undergo a distinct mode of exocytosis termed membrane crumpling, in which actomyosin-driven compression expels cargo while limiting complete integration of vesicle membrane into the apical surface, thereby preserving membrane homeostasis and cell morphology (Kamalesh et al., 2021)(Fig. 1A).

**Figure 1:**
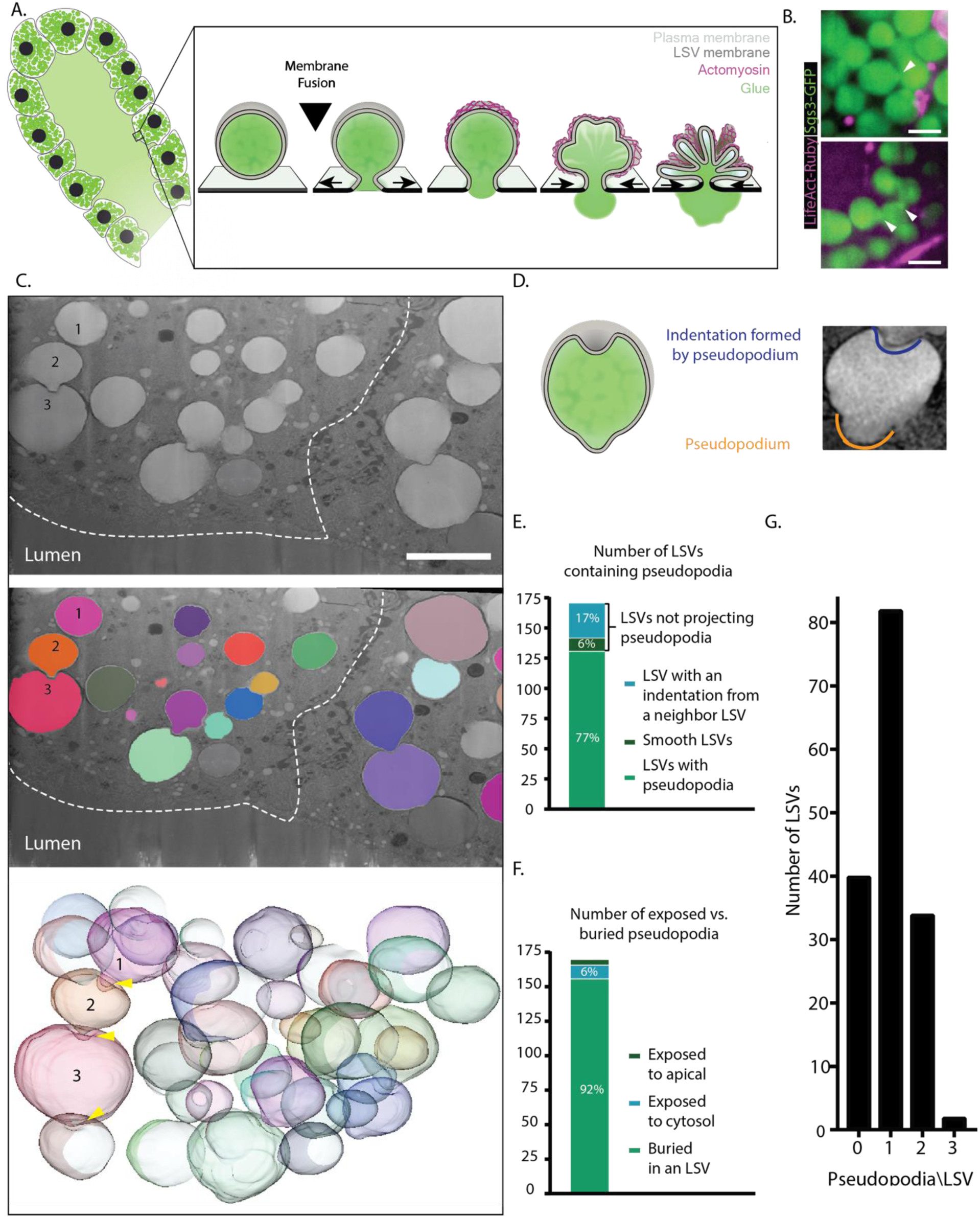
Vesicular pseudopodia are frequently found in *Drosophila* larval salivary glands. **A.** A schematic illustration of the process of exocytosis in the *Drosophil*a salivary gland. Upon fusion of the Glue-filled large secretory vesicles (LSV) with the apical membrane, an actomyosin coat is assembled around the LSV, mediating vesicular membrane crumpling and subsequent content release to the lumen. **B.** Vesicular pseudopodia as seen by confocal microscopy (white arrowheads). Though visible, these are challenging to capture in living samples. Scale bar: 5 μm. **C. Top:** Representative cross-section of a FIB-SEM depicting two secretory cells. The border of the left cell is defined by a white dashed line. LSVs numbered 1-3 are marked to enable tracking in all panels. Scale bar: 5 μm. **Middle:** Segmentation of the vesicles in the secretory cells. **Bottom:** Volume rendering of the segmented surface of the LSVs. **D.** Left: an illustration of an LSV in cross-section projecting one pseudopodium and displaying a dent formed by a pseudopodium projected from a neighboring vesicle. Right: depiction of a cropped LSV from a FIB-SEM cross-section with a pseudopodium (orange) and a dent (blue). **E.** Quantification of the population of LSVs: 77% of LSVs project at least one pseudopodium to their neighbors, 17% have only an indentation which is formed by the pseudopodium of another LSV, and the remaining 6% are smooth (*N*_*larva*_=3, *n*_*LSVs*_=171). **F.** Among the observed pseudopodia, 92% are buried within an indentation of a neighboring LSV. The remaining pseudopodia are either exposed to the cytosol or exposed to the apical surface (6% and 2%, respectively; *N*_*larva*_=3, *n*_*pseudopodia*_=170). **G.** Quantification of the number of pseudopodia/LSV. Most LSVs project only one pseudopodium from their membrane, and rarely more than 2 pseudopodia are projected from a single LSV (*N*_*larva*_=3, *n*_*LSVs*_=158). Taken together, these results suggest an elaborate network that connects the LSVs within the larval salivary gland secretory cells.

Recent work further suggests that the LSV membrane is spatially patterned during secretion. The inverse-BAR domain protein Missing in Metastasis (MIM) localizes to the future fusion site before fusion and regulates fusion-pore dynamics, and structured recruitment of RhoGEF2 and myosin II generates anisotropic contractile organization on fused vesicles (Biton et al., 2023; Kamalesh et al., 2024). Together, these findings suggest that the site of fusion is specified before exocytosis begins. Here, we test the hypothesis that vesicular pseudopodia constitute a preorganized fusion domain on LSV membranes that defines the site of fusion with the apical membrane. Using *Drosophila* larval salivary glands, volume electron microscopy, live imaging, proteomics and three-dimensional correlative light and electron microscopy, we ask how pseudopodia are organized, whether they mark the site of membrane fusion, and which vesicular and apical membrane components contribute to this spatially defined exocytic interface.

## Results

### LSVs in *Drosophila* larval salivary glands are interconnected by vesicular pseudopodia

To assess the ultrastructure of LSVs and determine the prevalence of vesicular pseudopodia, we first imaged salivary glands co-expressing the cargo marker Sgs3-GFP (Costantino et al., 2008) and the F-actin marker LifeAct-Ruby. Although pseudopodia could occasionally be visualized by fluorescence microscopy, the high density of LSVs, together with the need to capture pseudopodia near their equatorial plane, prevented robust quantification by light microscopy (Fig. 1B). We therefore used focused ion beam scanning electron microscopy (FIB-SEM) tomography to characterize vesicular pseudopodia in three dimensions and at higher resolution.

FIB-SEM revealed frequent pseudopodia between neighboring LSVs. These structures typically projected from one LSV into a corresponding indentation on an adjacent LSV, creating an interconnected network of vesicles within the cell (Fig. 1C, D). To quantify their prevalence, we scored 171 LSVs from three salivary glands. Of these, 77% projected at least one pseudopodium, 17% did not project a pseudopodium but contained an indentation, and only 6% were smooth, with neither a pseudopodium nor an indentation (Fig. 1E, S1A). Most pseudopodia were buried within an indentation in a neighboring LSV (92%), whereas the remaining pseudopodia were exposed to the cytosol or near the apical membrane (Fig. 1F).

Most LSVs projected a single pseudopodium and contained a single indentation, whereas vesicles bearing more than two pseudopodia were rare (Fig. 1G, S1A). In addition, pseudopodia appeared to be polarized towards the apical surface (Fig. 1C, S1B-C, Supplementary Movie 1). Together, these observations show that vesicular pseudopodia are a prevalent and polarized structural feature of LSVs, forming an interconnected vesicle network oriented towards the apical membrane.

### Fusion with the apical membrane occurs at exposed vesicular pseudopodia

Having established that most LSVs are interconnected through vesicular pseudopodia, we next asked whether exposed pseudopodia participate in fusion with the apical membrane. In the FIB-SEM volumes, we observed LSVs positioned near the apical surface with exposed pseudopodia oriented towards the apical membrane (Fig. 2A). In some cases, these pseudopodia appeared continuous with narrow membrane necks connecting the LSV to the apical surface, consistent with fusion pores forming at exposed pseudopodia (Fig. 2B). These observations suggested that vesicular pseudopodia are not merely structural connections between neighboring LSVs, but may define the site where LSVs fuse with the apical membrane.

**Figure 2:**
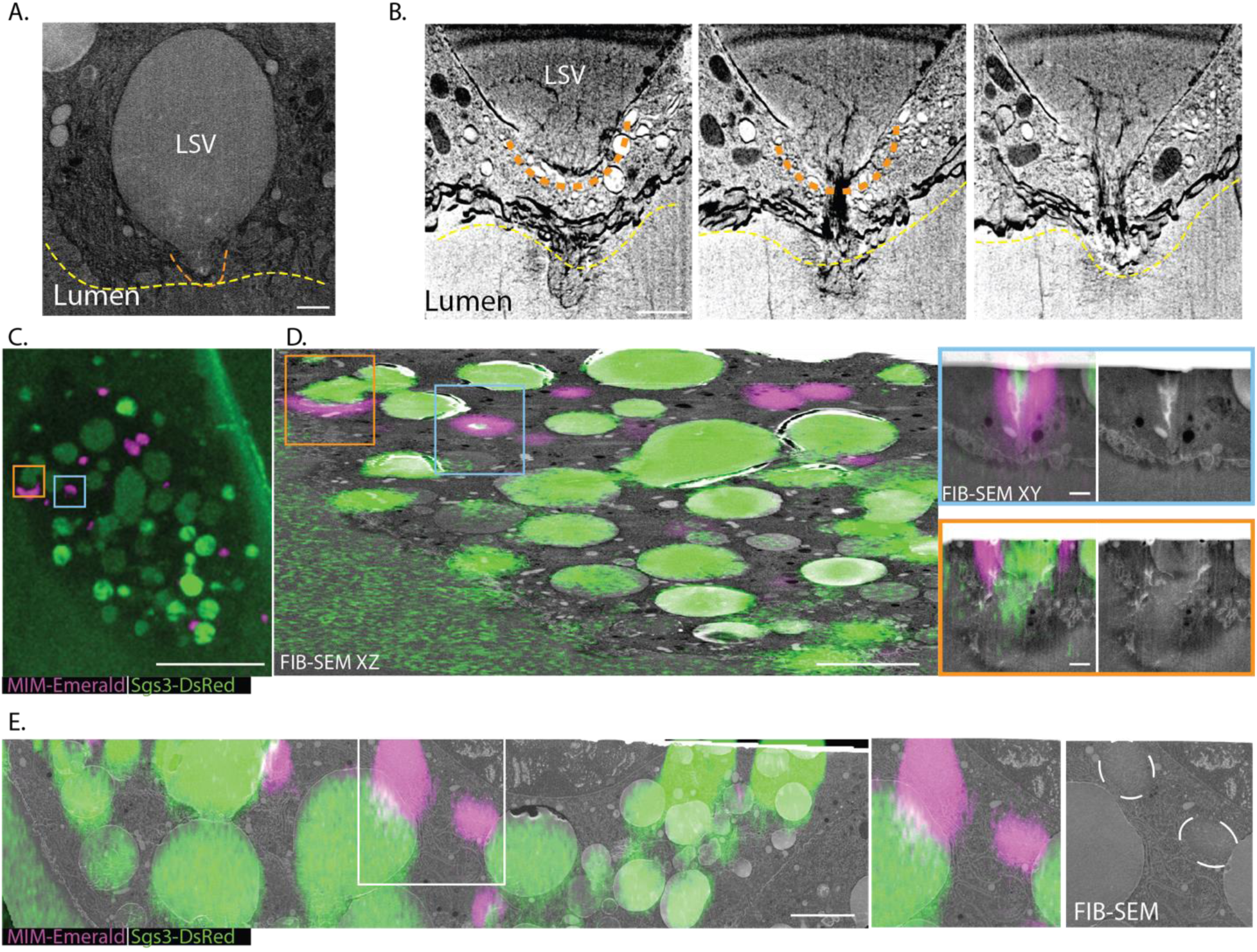
MIM-Emerald is localized to exposed pseudopodia where fusion happens. **A.** LSV with an exposed pseudopodium (dashed orange line) in close proximity to the apical surface (dashed yellow line). Scale bar 1 μm. **B.** Three slices through a FIB-SEM stack depicting a fused LSV with an extended protrusion similar to pseudopodia in shape. Scale bar: 1 μm. **C.** Optical slice through a resin-embedded sample depicting one secretory cell labeled with Sgs3-DsRed (green) and MIM-Emerald (magenta). Scale bar: 20 μm. **D.** Overlay of the transformed confocal stack of the cell in **A** with FIB-SEM tomography data. The correlation was performed using the LSVs as fiducial markers in both datasets. In occasions where MIM-Emerald signal surrounds LSVs (orange and teal squares), the underlying FIB-SEM data showed either an exposed and elongated pseudopodium (teal), or a fused LSVs (orange). Scale bars: 5 μm, inset 1 μm. **E. Left:** Overlay of transformed confocal stack and a FIB-SEM image of the same area in glands expressing Sgs3-DsRed (green) and MIM-Emerald (magenta). Scale bar 5 μm. **Right**: Enlarged part from the FIB-SEM cross-section on the **Left** (white rectangle) showing that MIM-Emerald puncta seem to concentrate in distinct areas in the cytosol.

We next sought molecular evidence that exposed pseudopodia correspond to the fusion site. We previously showed that the I-BAR domain protein Missing in Metastasis (MIM) localizes to the future fusion site on the vesicular membrane prior to fusion and subsequently remains associated with the fusion pore (Biton et al., 2023). Considering our current ultrastructural analysis, these earlier observations are consistent with MIM marking exposed pseudopodia positioned at the apical surface. To test this directly, we used three-dimensional correlative light and electron microscopy (3D-CLEM) to determine the localization of MIM relative to pseudopodia within whole secretory cells. We imaged glands co-expressing MIM-Emerald, a functional fluorescently tagged version of MIM, together with the glue cargo marker Sgs3-DsRed, which served as a fiducial marker for correlating fluorescence and electron microscopy volumes (Biton et al., 2023; Costantino et al., 2008).

3D-CLEM revealed that MIM-Emerald forms rings around exposed pseudopodia located close to the apical membrane and around LSVs that had already fused with the apical surface (Fig. 2C, D). In contrast, MIM-Emerald was not detected around buried pseudopodia positioned within indentations of neighboring LSVs. We also observed large cytosolic MIM-Emerald clusters with distinct electron density and texture compared with the surrounding cytoplasm (Fig. 2E), but these structures did not correspond to buried pseudopodia. Thus, we conclude that MIM is selectively associated with exposed pseudopodia near the apical membrane.

To determine whether MIM contributes to pseudopodia organization, we analyzed LSV morphology in MIM knockout glands. Loss of MIM did not grossly affect the frequency of pseudopodia, but pseudopodia displayed a reduced average aspect ratio compared with controls (Fig. S2). These results suggest that MIM is not required for pseudopodium formation per se, but may contribute to the elongation or stabilization of exposed pseudopodia at the apical surface. Together, the presence of fusion pores at exposed vesicular pseudopodia adjacent to the apical cell surface and the selective recruitment of MIM to these structures, support the conclusion that LSV fusion with the apical membrane occurs at exposed vesicular pseudopodia.

### MIM pull-down identifies the exocyst complex as a candidate pseudopodal component

Because MIM localizes to exposed pseudopodia near the apical membrane, we used it as a bait to identify additional proteins associated with the pseudopodal fusion domain. Since antibodies against endogenous *Drosophila* MIM are not commercially available, we performed immunoprecipitation from salivary glands expressing MIM-Emerald, using an antibody against the Emerald tag. Glands were dissected from third-instar larvae and treated ex vivo immediately after dissection with the cleavable, cell-permeable crosslinker dithiobis(succinimidyl propionate) (DSP), in order to preserve both stable and transient protein interactions. MIM-Emerald immunoprecipitation was then followed by mass spectrometry analysis (Fig. S3A). 746 proteins were significantly enriched in the MIM-Emerald samples compared with wild-type control glands (yw) (Fig. 3A, Supplementary Table 1), with MIM identified as the most enriched protein in the dataset, supporting the specificity of the approach. Among the hits were two previously reported MIM-interacting proteins, Cortactin and Downstream of receptor kinase (Drk). Cortactin, an Arp2/3 activator, directly binds MIM through an interaction between its SH3 domain and the MIM proline-rich domain (Lin et al., 2005; Quinones et al., 2010). MIM was also previously identified as a Drk-associated protein in a tandem affinity purification–mass spectrometry screen for Drk interactors (Friedman et al., 2011). Consistent with these known MIM interaction partners, proteins enriched in the MIM-Emerald pull-down were associated with biological processes related to actin organization, protein trafficking and endocytosis (Fig. 3B-C; Fig. S3B-C; Supplementary Table 1).

**Figure 3:**
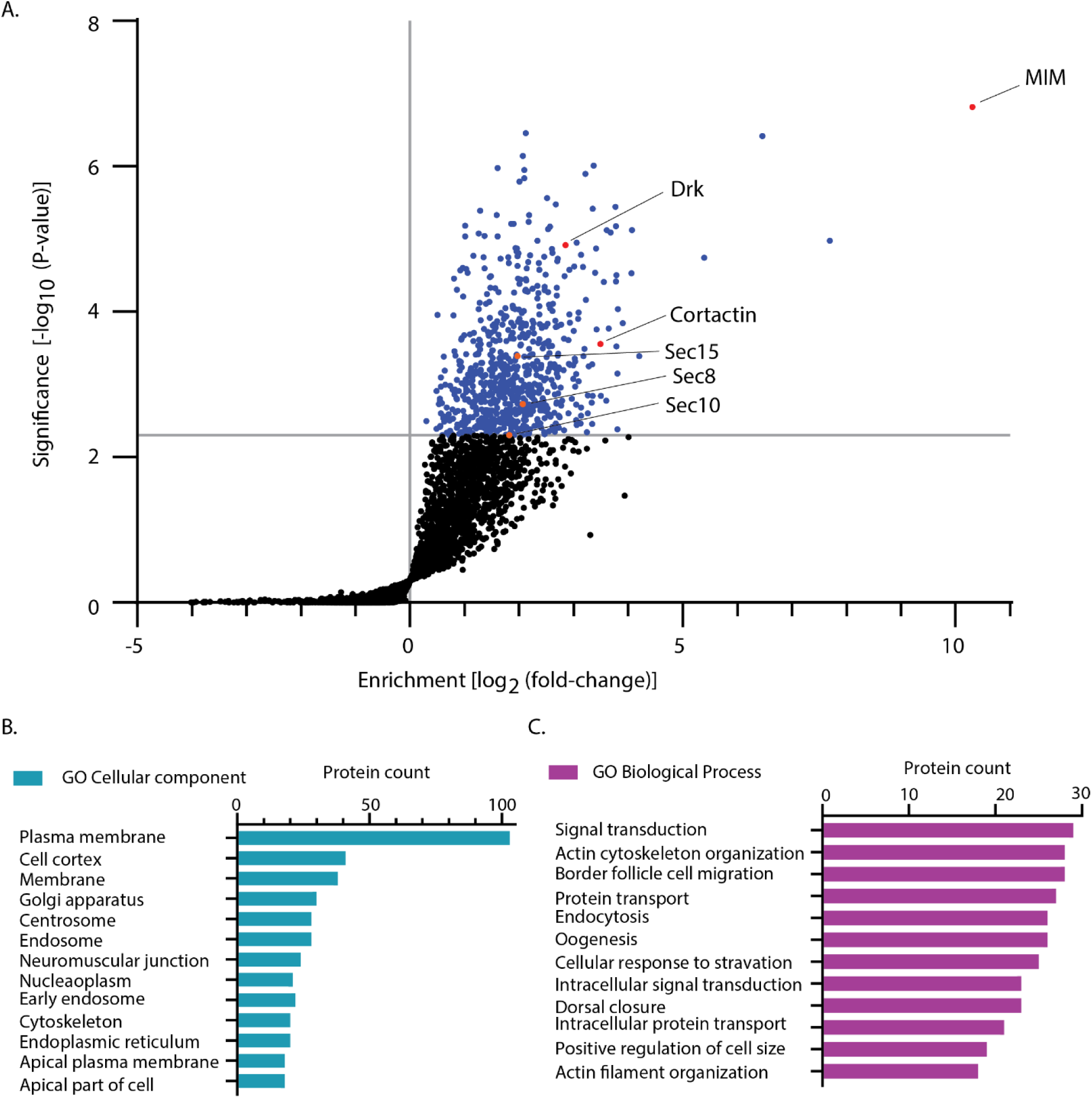
Exocyst subunits are enriched in a MIM pull-down **A.** Volcano plot of mass-spectrometry-based proteomics results from an analysis of immunoprecipitation using an antibody against Emerald-tagged MIM (in red), from lysates of crosslinked salivary glands. The results show significantly enriched proteins (in blue), identified using one-sided Student’s *t* test (FDR <0.01, S0 = 0.25). Among the identified proteins are Cortactin and downstream receptor kinase (Drk), both known direct interactors of MIM (in red). Three exocyst complex subunits (Sec8, Sec15 and sec10) were enriched in the MIM-Emerald samples (in orange). Volcano plot represents experiments performed in four biological replicates for each condition. **B.-C.** Gene ontology (GO) terms analysis of the significantly enriched proteins in the MIM-Emerald pull-down representing top hits in the GO cellular component (**B.**) and cellular component biological process (**C.**) categories.

Notably, three subunits of the exocyst complex were significantly enriched in the MIM-Emerald pull-down: Sec8, Sec15 and Sec10, corresponding to exocyst subunits 4, 6 and 5, respectively (Fig. 3A). The exocyst is an evolutionarily conserved octameric tethering complex that functions in targeted vesicle delivery and secretion, and was originally identified in budding yeast mutants defective in secretion (TerBush et al., 1996). In several systems, the exocyst participates in vesicle tethering to the plasma membrane and can interact with SNARE-associated machinery to promote membrane fusion (An et al., 2021). The identification of Sec15 was of particular interest, because Sec15 is a known Rab11 effector in exocytic trafficking (Wu et al., 2005; Zhang et al., 2004), and Rab11 localizes to secretory granule membranes and is required for normal exocytosis in *Drosophila* larval salivary glands (Neuman et al., 2021). We therefore hypothesized that Sec15 may associate with LSVs and localize to vesicular pseudopodia during secretion.

### The exocyst subunit Sec15 localizes to the site of fusion and to vesicular pseudopodia

To determine Sec15 localization during secretion, we imaged secreting larval salivary glands expressing Sgs3-DsRed with GFP-tagged Sec15 (GFP-Sec15) over-expressed via the C135-GAL4 driver, at different stages of the secretion process. In early-secreting salivary glands, GFP-Sec15 puncta were primarily observed between the LSVs, with minimal localization on the apical membrane (Fig. 4A-B). Immunofluorescence in fixed salivary glands and at native expression levels also confirmed Sec15 localization to the interface between vesicles (Fig. S4). In addition to this localization, in mid- and late-secreting glands, GFP-Sec15 formed large clusters on the apical membrane (Fig. 4C-D). The change in GFP-Sec15 localization suggested that Sec15 may localize to vesicular pseudopodia and accumulate at the apical membrane as secretion progresses.

**Figure 4:**
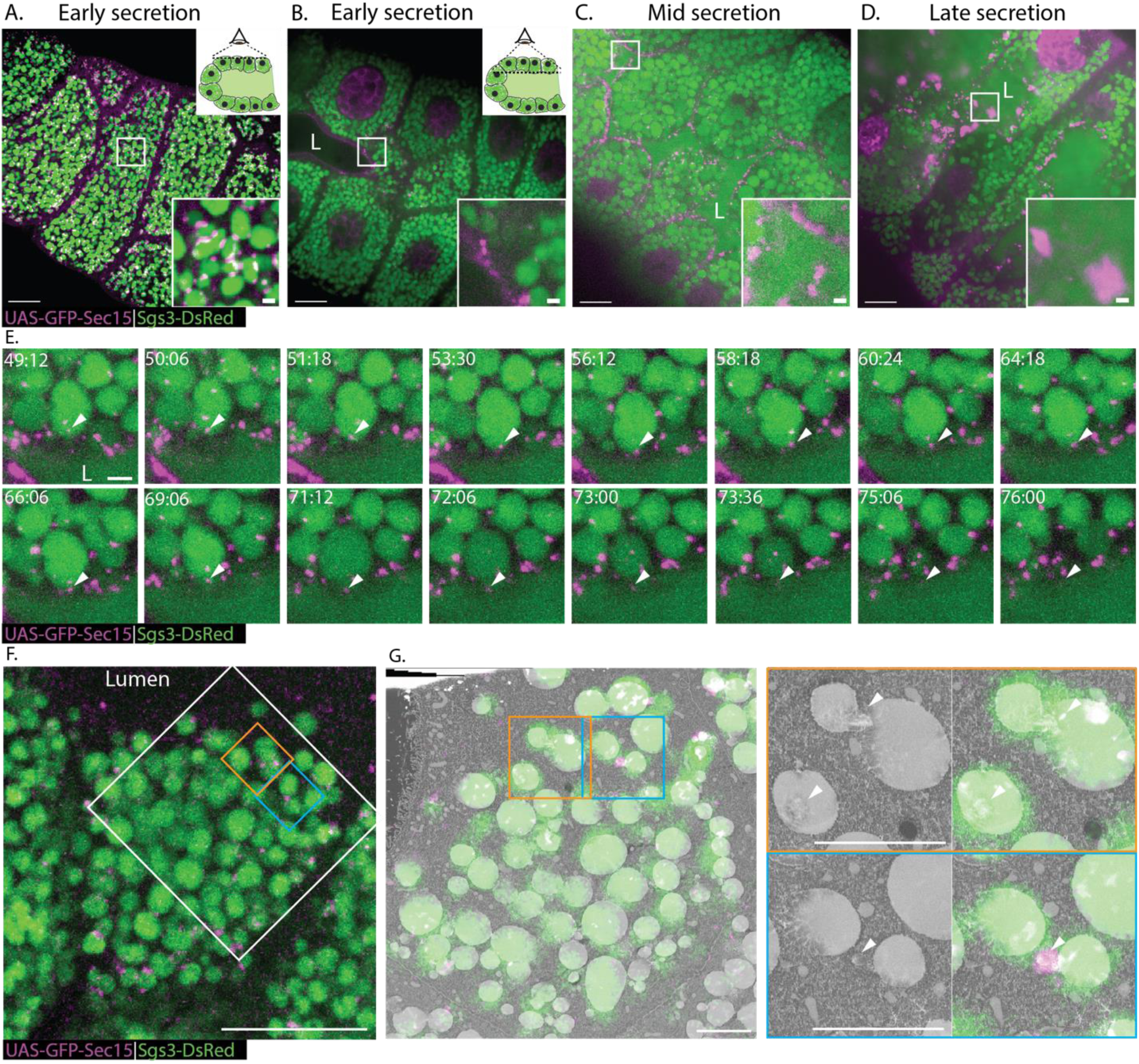
The exocyst subunit Sec15 localizes to vesicular pseudopodia. **A.** Representative images from confocal live imaging of glands expressing GFP-Sec15 under the UAS expression and Sgs3-DsRed. During the early stages of secretion GFP-Sec15 localizes between the LSVs and **B.** in small puncta at the apical membrane. **C.-D.** As secretion propagates, GFP-Sec15 puncta accumulate at the apical membrane and increases in size, as it accumulates there. Scale bars: 20μm, inset 2μm. **E.** Confocal timelapse series of an LSV fusing with the apical surface. GFP-Sec15 puncta (white arrowhead) are present during secretion at the interface between the LSV and the apical membrane. Time mm:ss; Scale bar: 2 μm. **F.** Optical slice through the resin-embedded sample depicting the same markers shown in **A-E**. The large bounding box relates to the area acquired with FIB-SEM tomography. Scale bar: 20 μm. **G.** Overlay of the transformed confocal stack of the cell in **F** with FIB-SEM tomography data. The correlation was performed using the LSVs as fiducial markers in both datasets. The two insets represent two localizations of GFP-Sec15. The first is to the vesicular pseudopodia (orange) and the second is to a vesicle that might be an endosome (teal). Scale bar: 5 μm. These results show that Sec15 is involved in the fusion of LSVs and is localized to the pseudopodia membrane domain. This further supports the hypothesis that vesicular pseudopodia are fusion membrane domains on the vesicular membrane.

To determine whether Sec15 localizes to LSVs throughout secretion, we followed GFP-Sec15 localization on LSVs by live-imaging. We observed that GFP-Sec15 puncta persist on LSVs close to the apical membrane. Notably, Sec15 puncta were positioned at sites where the LSVs fused with the apical membrane, and persisted post-fusion (Fig. 4E, Supplementary Movie 2). This suggests that Sec15 indeed localizes to the LSV membrane before fusion and persists during fusion and cargo release, strengthening the hypothesis that it localizes to pseudopodia.

To determine whether Sec15 localizes specifically to pseudopodia, we performed 3D-CLEM of salivary glands expressing Sgs3-DsRed and GFP-Sec15. The correlations showed that Sec15-GFP localizes to pseudopodia (Fig. 4F-G). Taken together, our results demonstrate Sec15 localizes to vesicular pseudopodia, likely to coordinate the tethering of both the vesicle and cell membranes before fusion.

### Tsp42Ee defines a fusion domain on the apical surface

Since Tetraspanins (TSPs) have been linked to SNARE-mediated fusion by organizing and remodeling membranes (Charrin et al., 2014; Dahmane et al., 2019; Dharan and Sorkin, 2024; Dharan et al., 2022; Earnest et al., 2017; Runge et al., 2007; van Deventer et al., 2021), we wondered whether they have a role in this process. To test the role of TSPs in LSV secretion, we first overexpressed the exogenous murine TSP CD63-GFP, previously shown to interfere with glue vesicle maturation, which occurs via homotypic fusion of smaller nascent vesicles (Ma et al., 2020). Overexpression of CD63-GFP resulted in large and deformed LSVs decorated with CD63-GFP on their membranes, as previously shown (Ma et al., 2020) (Fig. 5A). Similar results were obtained upon overexpression of untagged *Drosophila* TSPs (Fig. S5A-B). Strikingly, live imaging of secreting glands revealed that as secretion propagates, glands became full of enormous over-sized vesicles connected to the lumen via large fusion stalks (Fig. 5A-B). These observations suggested that upon the initiation of secretion, mature glue vesicles begin to fuse with one another in a manner reminiscent of compound exocytosis. To test this hypothesis, we imaged glands overexpressing CD63-GFP together with Sgs3-GFP and LifeAct-Ruby to label the cargo and F-actin, respectively. Time-lapse microscopy showed that individual LSVs indeed fuse with oversized LSVs in a compound exocytosis-like manner. Compound exocytosis events captured via time-lapse imaging suggested that the LSV-LSV fusions initiated only after an initial fusion event between an LSV and the apical membrane and the onset of secretion had occurred (Fig. 5B). To test whether CD63-GFP is enriched in the pseudopodia, we used 3D-CLEM (Fig. S5C). However, we could not conclude that CD63-GFP corresponded with areas where pseudopodia were visible (Fig. S5C). These findings, taken together, show that TSP overexpression perturbs the fusion process, leading to oversized LSVs.

**Figure 5:**
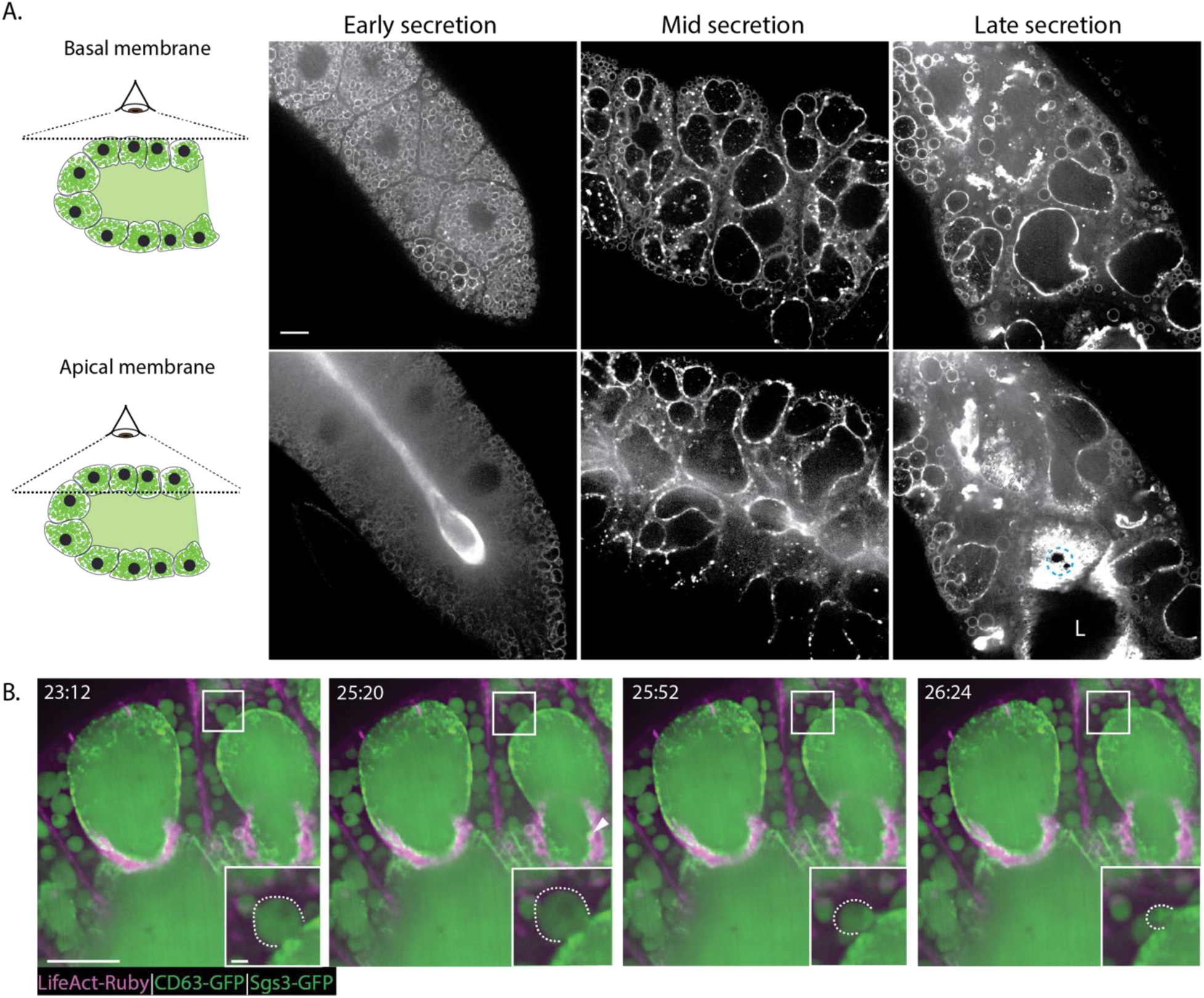
Over-expression of the mammalian TSPs CD63 leads to LSV deformation and compound exocytosis. **A.** Over-expression of the murine exogenous TSP CD63-GFP using UAS-based expression driven by C135-GAL4. The top panels represent optical slices closer to the basal surface of the salivary gland, while the bottom panels represent optical slices close to, or at the border of the apical cell surface and the lumen of the salivary gland. At earlier stages, the vesicles are round and similar in size to regular LSVs. As secretion propagates, LSVs fuse one with each other to form larger LSVs that are fused to the apical membrane (Blue dashed circle). L – lumen, scale bar: 20 μm. **B.** Confocal time-lapse series from a gland expressing UAS-CD63-GFP (green; vesicle membrane), Sgs3-GFP (green; vesicle lumen) and UAS LifeAct Ruby (red). The bottom inset shows a larger magnification of an LSV (dashed line) fusing via apparent compound exocytosis with an over-sized LSV which has fused with the apical membrane. Time mm:ss; L – lumen, scale bars: 20 µm, insets 5 µm.

The highest natively expressed TSP in the salivary gland is Tsp42Ee (Consortium et al., 2011; Li et al., 2014) (Fig S6A). To follow the localization of Tsp42Ee in secretory cells, we imaged glands expressing Tsp42Ee-EGFP under its native promoter, together with myristilated RFP (Myr-RFP), which labels the apical and vesicular membranes. We observed that Tsp42Ee puncta were localized at the apical membrane and in the cytosol (Fig. 6A). To test whether Tsp42Ee has a role in exocytosis, we performed RNAi knockdown experiments. Knocking down Tsp42Ee resulted in decreased frequency of “membrane crumpling” exocytosis events (32±3% and 26±5%) compared to wild-type (60±3%), and an increase in vesicle stalling and full collapse (Fig. 6B), similar to MIM knockdown (Biton et al., 2023). These results imply that Tsp42Ee contributes to the fusion process and therefore may localize to fusion sites.

**Figure 6:**
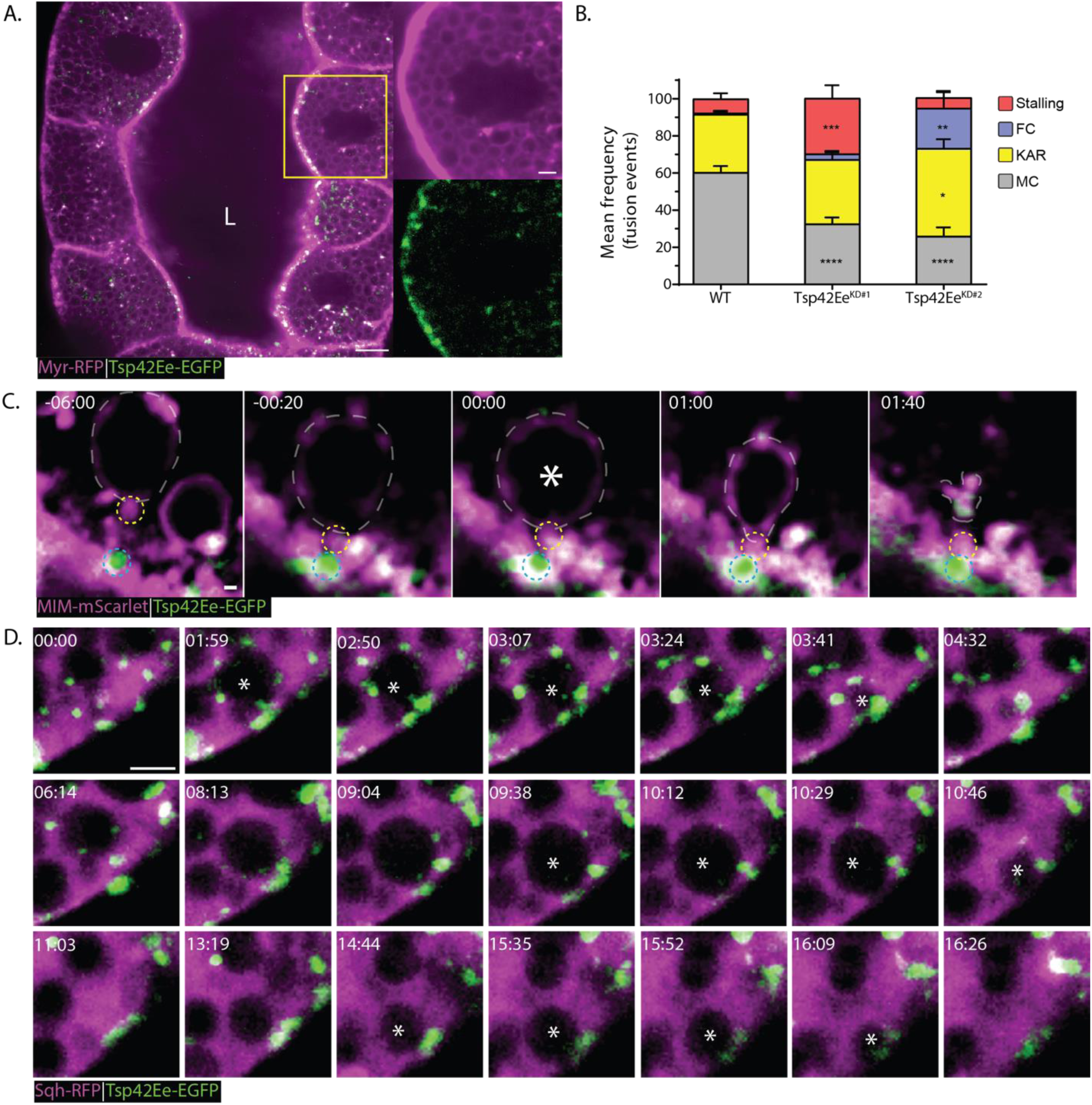
Tsp42Ee defines fusion domains on the apical surface. **A.** Low magnification of a salivary gland expressing the membrane marker Myr-RFP (magenta) and Tsp42Ee-EGFP (green). Tsp42Ee is found predominantly on the apical membrane in puncta and in between LSVs. L-lumen, scale bars: 20 µm, inset: 5 µm. **B**. Mean frequency (%) of stalling, full collapse (FC), kiss-and-run (KAR), and membrane crumpling (MC) in WT and Tsp42Ee KD #1 and #2 salivary glands (expressing Tsp42Ee-RNAi constructs BDSC_62303 & BDSC_77368, respectively) using the Sgs3-GFP and LifeAct-Ruby markers. The majority of fusion events observed in WT salivary glands with Sgs3-GFP and LifeAct-Ruby markers result in membrane crumpling events (60 ± 3%), while in both KD#1 and KD#2 a lower frequency of membrane crumpling events was observed (32 ± 3%****; 26 ± 5%**** respectively). Under KD #1, a significantly higher frequency of stalling events was observed (8 ± 3% in WT; 30 ± 7%*** in KD#1), while under KD #2 a significantly higher frequency of KAR and FC events were observed (23 ± 5%; 1 ± 1% in WT respectively; 32 ± 15%*; 21 ± 16%** in KD#2) N (glands) = 3, n (events) ≥ 130. Two-tailed unpaired multiple t-tests corrected using the Holm-Sidak method. UAS-based expression is driven by C135-GAL4. **C**. Time-lapse series (presented as confocal intensity-projections), of representative LSVs from salivary glands expressing both the MIM-mScarlet (Magenta; under UAS control) and the Tsp42Ee-EGFP (Green; under the endogenous promotor; BDSC_51588) markers. Fusion onset (marked by vesicle swelling) - white asterisk, LSV - White dashed transparent polygon line (marked using the MIM-mScarlet channel), MIM puncta at fusion site (LSV) - Yellow dashed circle, Tsp42Ee puncta at fusion site (apical) - blue dashed circle. Time mm:ss; relative to fusion. Scale bar: 1 µm. **D**. Salivary gland expressing Sqh-mCherry (magenta) and Tsp42Ee-EGFP (green) under their endogenous promoters. LSVs can be seen by the shaded areas in Sqh-mCherry signal. Time-lapse series of a small area from a secreting cell. showing that four vesicles fuse to the same area on the apical membrane (fusing vesicles marked in asterisk) within less than 20 minutes. Time mm:ss; Scale bar: 5 µm.

To assess Tsp42Ee localization to fusion sites, we co-expressed Tsp42Ee-EGFP and MIM-mScarlet, which localizes to the sites of fusion before the onset of membrane fusion and remains at the fusion pore throughout secretion (Biton et al., 2023). Strikingly, live imaging showed that MIM-mScarlet and Tsp42Ee-EGFP fluorescence appear clearly separate prior to fusion, but co-localize once fusion is initiated (Fig. 6C, S6D, Supplementary Movie 3). Moreover, we found that multiple fusion events occur in sequence at sites marked by Tsps42E-GFP at the apical surface (Fig. 6D, S6B; Supplementary Movies 4-5). Unfortunately, we could not precisely localize Tsps42Ee-EGFP using CLEM because the fluorescence was not preserved in resin, likely due to endogenous expression levels. Interestingly, Tsp42Ee-EGFP localizes to the surface of oversized vesicles in glands overexpressing CD63-mCherry, instead of to the apical surface (Fig. S6C). Taken together, these results are consistent with a model in which Tsp42Ee is a component of fusion domains on the apical membrane, where multiple vesicles fuse in sequence. Partial loss of Tsp42Ee perturbs membrane crumpling exocytosis, demonstrating that it is an essential regulator of this process. Moreover, when Tsp42Ee localization shifts to the vesicular membrane, vesicles now fuse with one another, resulting in compound exocytosis.

## Discussion

Large secretory vesicles (LSVs) pose a fundamental scaling problem for regulated exocytosis. Their micron-scale dimensions raise a central question of how LSVs spatially organize their fusion machinery, so as to ensure efficient and timely fusion with the apical membrane. In this study, we show that vesicular pseudopodia provide a solution to this problem in the *Drosophila* larval salivary gland. Using FIB-SEM, live imaging, proteomics and 3D-CLEM, we find that most LSVs are interconnected through pseudopodia that appear to form a polarized network oriented towards the apical surface. Exposed pseudopodia are found at the apical membrane and are associated with narrow fusion pores, indicating that fusion occurs at these structures. We further identify MIM and the exocyst subunit Sec15 as components associated with the vesicular pseudopodal fusion domain, and show that the tetraspanin Tsp42Ee marks a complementary fusion-prone domain on the apical membrane. Together, our findings support a model in which LSV fusion is not initiated randomly along the vesicle surface, but is spatially defined by the alignment of two specialized membrane domains: a vesicular pseudopodium and an apical tetraspanin-rich membrane domain.

The idea that vesicular pseudopodia participate in secretion was proposed more than five decades ago, based on ultrastructural observations in mammalian salivary glands. Early work described protrusions extending from secretory granules and suggested that they were oriented towards the apical membrane and might represent the site of fusion (Odajima and Nakane, 1984; Schramm et al., 1972; Selinger et al., 1974). Similar structures were later observed across diverse secretory systems, including salivary glands, lacrimal glands, mammary glands, nasal turbinate glands, insect secretory tissues and other specialized secretory epithelia (Scher and Avinoam, 2025; Tandler and Phillips, 1993; Tandler et al., 2000; Wu et al., 2006). However, because most of these observations relied on static two-dimensional electron micrographs, the prevalence, organization and molecular identity of pseudopodia remained unknown. Our volume EM analysis now shows that pseudopodia are not rare or incidental structures in *Drosophila* salivary glands. Rather, they are a prevalent feature of mature LSVs, with the vast majority of vesicles projecting at least one pseudopodium and containing a corresponding indentation.

Our observations directly support the idea that pseudopodia define the vesicular fusion site. We observed exposed pseudopodia at the apical surface, including examples in which a narrow neck connects the vesicle to the apical membrane, consistent with a fusion pore forming at the pseudopodium. These ultrastructural observations provide an explanation for our previous finding that MIM localizes to the future fusion site before membrane fusion and remains associated with the fusion pore during secretion (Biton et al., 2023). At the time, this localization was interpreted as marking a vesicular site that would eventually fuse with the apical membrane. The present 3D-CLEM data now allow us to assign this site to an exposed vesicular pseudopodium.

MIM may also contribute actively to pseudopodium architecture. MIM is an inverse-BAR domain protein capable of sensing and generating curved membrane protrusions (Ahmed et al., 2010; Mattila et al., 2007; Nishimura et al., 2021; Tsai et al., 2022). In MIM mutant glands, pseudopodia were still present, indicating that MIM is not strictly required for pseudopodium formation. However, their reduced aspect ratio (Fig. S2) suggests that MIM may promote pseudopodium elongation, stabilization, or remodeling once pseudopodia become exposed to the cytosol (Fig. 7). Such a model fits well with the selective localization of MIM to exposed pseudopodia rather than buried pseudopodia. Buried pseudopodia are enclosed within indentations of neighboring vesicles and may be inaccessible to cytosolic MIM, whereas exposed pseudopodia present a curved membrane surface to the cytosol.

**Figure 7:**
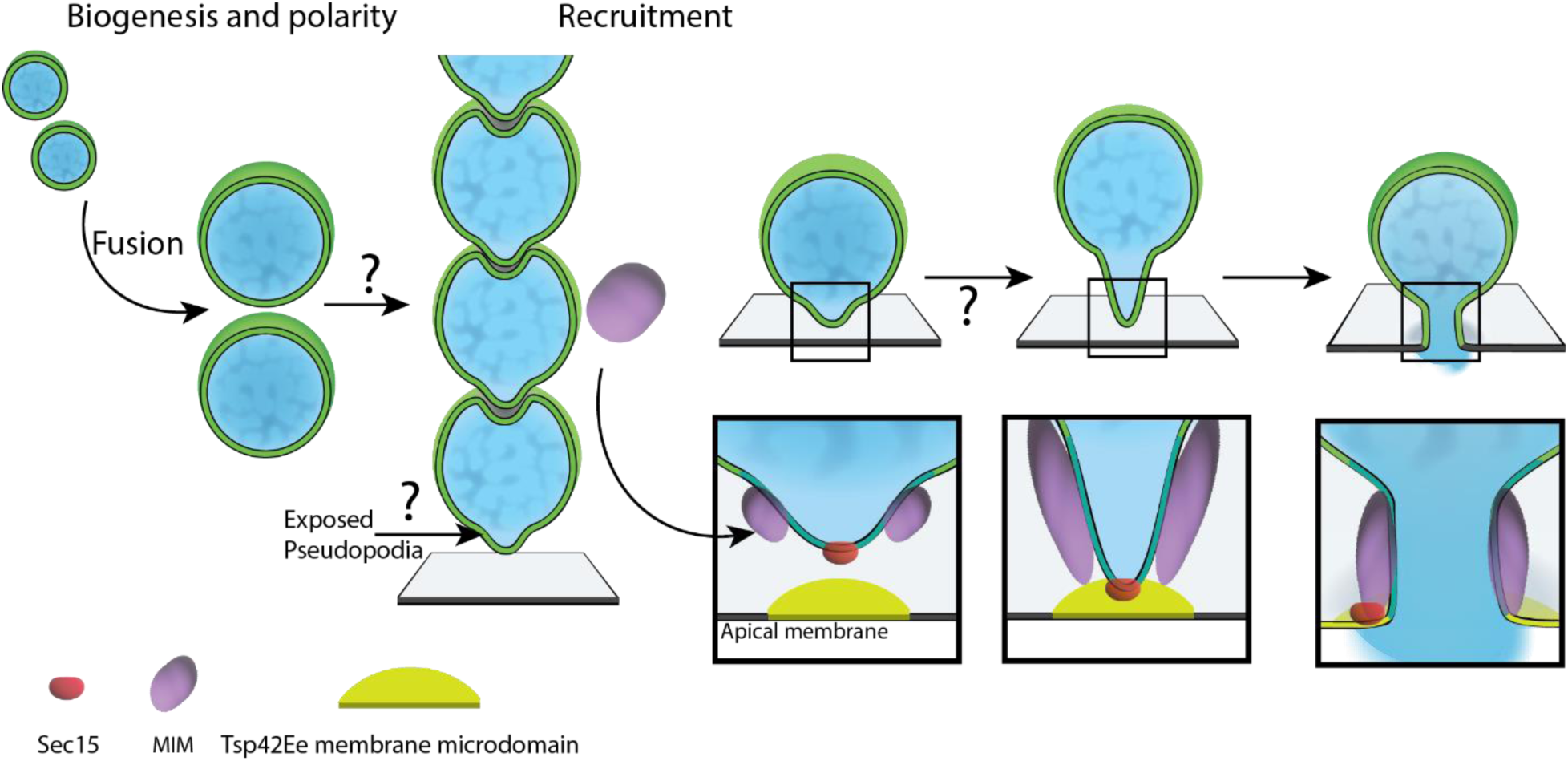
Schematic model depicting a role for pseudopodia as a vesicular fusion domain. During the process of LSVs maturation, secretory vesicles fuse to one another until reaching their full size. How and when pseudopodia are formed during the LSVs maturation process remains to be determined. However, in the mature salivary gland, the LSVs form a polarized network throughout the secretory cells, oriented towards the apical membrane. At this point, the exocyst subunit Sec15 is already localized to the vesicular pseudopodia. On the opposite membrane, the tetraspanin Tsp42Ee defines a fusion domain on the apical surface. Once secretion initiates, MIM localizes to the newly exposed pseudopodium close to the apical surface. The pseudopodium then elongates towards the apical membrane where the exocyst complex tethers both membrane domains, leading to membrane fusion and membrane crumpling exocytosis.

The MIM pull-down results further support the idea that exposed pseudopodia are molecularly specialized. Among the enriched proteins were three subunits of the conserved exocyst tethering complex: Sec8, Sec15, and Sec10 (An et al., 2021; TerBush et al., 1996). Sec15 is a known Rab11 effector in exocytic trafficking (Wu et al., 2005; Zhang et al., 2004), and Rab11 was previously shown to localize to secretory granule membranes and to be required for normal secretion in *Drosophila* salivary glands (Neuman et al., 2021). Our live-imaging and 3D-CLEM data show that Sec15 localizes to LSVs, persists at sites of fusion, and is present at pseudopodia (Fig. 4). These findings reinforce the function of pseudopodia as a platform for exocyst-mediated tethering and subsequent fusion of LSVs to the apical membrane.

This view aligns with recent work showing that the exocyst regulates multiple stages of LSV maturation and secretion (Freire et al., 2024; Lee et al., 2025). LSVs mature through homotypic fusion, a process that increases vesicle size before the ecdysone-triggered wave of secretion (Biyasheva et al., 2001; Burgess et al., 2011; Ma et al., 2020). The exocyst, therefore, appears to operate at more than one stage of the LSV lifecycle, contributing to homotypic fusion events that generate mature LSVs and to tethering and fusion to the apical membrane. This dual role raises an exciting possibility that homotypic fusion of LSV precursors arrests once they become secretion competent by spatially restricting exocyst activity to tethering, while allowing fusion only when the pseudopodium is exposed and properly aligned with an apical membrane domain (Fig. 7).

On the vesicular side, the pseudopodium concentrates or recruits MIM, Sec15 and likely additional fusion regulators. On the target membrane side, our data identify Tsp42Ee as a functional component of apical fusion domains, consistent with the role of tetraspanins in organizing membrane microdomains and regulating the local availability of partner proteins, including SNAREs (Charrin et al., 2014; Dharan and Sorkin, 2024; Dharan et al., 2022; van Deventer et al., 2021; Wojnacki et al., 2023). We show that in the salivary gland, Tsp42Ee localizes predominantly to the apical membrane, and its knockdown decreases the frequency of membrane crumpling events while increasing abnormal vesicle stalling and full-collapse-like behavior (Fig. 6). Moreover, MIM and Tsp42Ee occupy opposing membranes before fusion and become coincident only after fusion has occurred (Fig. 6, S6). This behavior is consistent with a model in which the MIM-positive pseudopodal domain on the vesicle engages a Tsp42Ee-positive domain on the apical membrane (Fig. 7). The apical domain may have several related functions. First, it may help define where LSVs fuse by locally organizing SNAREs, lipids, tethers, or other fusion regulators at the apical membrane. Second, it may help maintain the identity of the fusion site after fusion by restricting diffusion between the apical membrane and the vesicle membrane (Kamalesh et al., 2021). Third, it may contribute to regulation of fusion-pore dynamics and thereby influence the mode of exocytosis. We previously showed that membrane crumpling requires a fusion pore that expands, stabilizes and subsequently constricts during actomyosin-mediated cargo release, and that disruption of this sequence redirects LSVs towards alternative modes: pores that fail to expand or reseal prematurely undergo kiss-and-run-like events, whereas pores that expand without stabilizing proceed to full collapse (Biton et al., 2023). The increased occurrence of stalling and kiss-and-run-like events following Tsp42Ee depletion therefore suggests that Tsp42Ee is required for productive pore expansion, enabling LSVs to proceed through membrane-crumpling exocytosis. Taken together, these observations suggest that fusion-pore regulation depends on coordinated organization of both the vesicular and apical membrane domains.

The compound exocytosis-like phenotype caused by tetraspanin perturbation further extends this model. Overexpression of murine CD63-GFP, as well as of *Drosophila* native tetraspanins, caused enlarged and deformed LSVs, consistent with previous work showing that CD63 overexpression affects homotypic vesicle fusion during glue-granule maturation (Ma et al., 2020). During secretion, CD63-GFP-overexpressing glands developed oversized vesicles connected to the lumen through large fusion stalks. Live imaging showed that under these conditions, Tsp42Ee is no longer restricted to the apical domain and that additional LSVs fuse with these enlarged vesicles in a compound exocytosis-like manner (Fig. 5, S6D). We speculate that disruption of the normal diffusion barrier at the fusion interface allows apical fusion-domain identity, including Tsp42Ee, to spread onto the membrane of a fused LSV. This would convert the fused vesicle into an ectopic target membrane for subsequent fusion events, thereby promoting compound exocytosis. Such a mechanism is consistent with our previous proposal that restricted mixing between vesicular and apical membranes is essential for maintaining membrane-crumpling exocytosis (Kamalesh et al., 2021; Biton et al., 2023).

In summary, the emerging model suggests that during larval development, glue proteins are packaged into secretory vesicles that mature through homotypic fusion (Biyasheva et al., 2001; Burgess et al., 2011; Ma et al., 2020). As vesicles mature into LSVs, our study reveals that they customarily form pseudopodia and corresponding indentations, creating a connected, polarized vesicle network. When secretion initiates, and vesicles begin to fuse, the pseudopodia become exposed near the apical surface. Upon exposure, the pseudopodal membrane that already contains the exocyst tethering complex, recruits MIM, which may elongate or stabilize the pseudopodium and contribute to fusion-pore regulation (Biton et al., 2023). Fusion then occurs between the pseudopodium and a Tsp42Ee-positive apical membrane domain (Fig. 7). After pore formation, the pore expands and stabilizes, allowing actomyosin recruitment to the fused vesicle membrane, which generates anisotropic contractile forces that drive vesicle crumpling and cargo expulsion while limiting incorporation of the vesicle membrane into the apical surface (Biton et al., 2023; Kamalesh et al., 2021; Kamalesh et al., 2024).

Several questions remain open. First, it is not yet clear when pseudopodia form. They may arise during homotypic fusion and vesicle maturation, or they may emerge later as mature LSVs become secretion competent. Second, the molecular mechanism that specifies pseudopodium polarity remains unknown. Our data show that pseudopodia are oriented towards the apical membrane and that MIM may contribute to their morphology, but whether polarity is imposed by apical cues, vesicle–vesicle packing, cytoskeletal organization, Rab-dependent membrane identity, or exocyst-mediated tethering remains to be determined. Third, the composition of the buried pseudopodium and the corresponding indentation is mostly unknown. Sec15 localizes to buried pseudopodia, while MIM appears to be selectively recruited to exposed pseudopodia, but surely other distinct proteins or lipids that organize vesicle-vesicle interactions are present. Finally, it is still unclear how diffusion becomes restricted after vesicle fusion. TSPs regulate both homotypic vesicle-vesicle and heterotypic vesicle-plasma membrane fusion, making them attractive candidates as organizers of these fusion domains; however, additional lipids, SNARE regulators, cortical actin, septins, or tethering factors may also contribute to maintaining a boundary between vesicular and apical membrane identities.

The cytosolic MIM-Emerald clusters observed by 3D-CLEM and live imaging may point to an additional layer of regulation. These clusters did not correspond to buried pseudopodia, but their distinct texture together with the observed enrichment of RNA-associated proteins in the MIM pull-down experiments suggest that MIM is stored or regulated in cytosolic assemblies before recruitment to exposed pseudopodia. At present, this remains speculative. It will be important to determine whether these MIM-positive clusters behave as condensates.

Overall, our findings provide direct evidence that vesicular pseudopodia define the site of LSV fusion with the apical membrane. They also identify molecular components associated with the opposing vesicular and apical membrane domains that mediate this process. By connecting earlier ultrastructural observations with modern live imaging, proteomics and 3D-CLEM, this work revives and extends the hypothesis that large secretory vesicles use specialized protrusive membrane domains to organize efficient, directional and spatially restricted secretion. In the *Drosophila* salivary gland, this organization allows the cell to coordinate vesicle maturation, apical targeting, regulation of fusion pore dynamics, cargo release, and maintenance of membrane homeostasis during a massive burst of secretion. More broadly, vesicular pseudopodia may represent a general adaptation of secretory tissues that rely on micron-scale vesicles, providing a structural and molecular solution to the challenges imposed by large-vesicle exocytosis.

## Materials and methods

### *Drosophila* strains and rearing conditions

Flies were obtained from the Bloomington *Drosophila* Stock Center (NIH P40OD018537), Zurich ORFeome Project (FlyORF; https://flyorf.ch/), and Vienna Drosophila Resource Center (VDRC; https://shop.vbc.ac.at/vdrc_store/). Sgs3-DsRed and UAS-Tsp29Fa were a kind gift from Jullie Brill (University of Toronto). MIMnull (Quinones et al., 2010) was a kind gift from Helen Zenner (University of Cambridge). UASp-CD63-GFP was a kind gift from Eli Arama (Weizmann Institute of Science). All fly lines used in this study are summarized in Supplementary Table 2.

Fly stocks were kept in flasks with standard fly food (cornmeal, molasses, and yeast media) in a temperature-controlled room at 21°C. For experimental procedures, fly lines were grown at 25°C without illumination. Lines for experiments were kept at low density (20-30 flies/bottle) and were moved to fresh bottles every 3-5 days. Larvae of both sexes were used interchangeably in this work because the salivary glands’ secretion process has no noticeable differences between males and females.

### Culturing larval salivary glands for live imaging

*Drosophila* 3^rd^ instar larvae were dissected in a watch glass filled with Schneider’s insect medium (SIM; Thermo Fisher Scientific) at room temperature (RT). Induction of secretion with the hormone Ecdysone was not used during this work and to image glands during secretion, glands with a swollen lumen were identified under an Olympus MVX10 stereoscope in the whole larva. Dissected glands were transferred to a 10 mm glass-bottom well in a 35 mm imaging plate (D35-10-0-N; Cellvis) filled with 100 µL of SIM.

### Confocal imaging

Samples were imaged using an Olympus IX83 inverted microscope with a Yokogawa automatic spinning disk confocal scanning unit (CSU-W1-T2). Images were acquired using a PLAPON 60×1.4 NA oil immersion objective. Signal detection was done using dual back-illuminated Prime 95B sCMOS cameras (Photometrics) controlled via VisiView software (Visitron Systems). Fluorescence excitation was applied using solid-state laser diodes (Toptica) at 488nm, 561nm and 647nm wavelengths and via 525/550 nm, 609/654 nm or 700/775 nm emission filters, respectively. For time-lapse imaging, dual cameras were used with an additional 561 long-pass D2 dichroic mirrors. All fluorescent micrographs were acquired at RT.

### Vesicle size quantification

Three random optical slices were taken from each confocal stack for vesicle cross-sectional area calculation. Each slice was processed using built-in Fiji functions (Schindelin et al., 2012). The slices were duplicated and two median filters (with 2 pixels and 50 pixels) radii were applied on the different copies of the same slice. Then the 50 pixels median filter image was subtracted from the 2 pixels median filter image for background noise removal. The resulting image was processed by contrast-limited adaptive histogram equalization (CLAHE) to enhance local contrast before a binary image was made using the threshold tool. The binary image was then processed by watershed to separate close objects. Analyze particles algorithm was applied to each binary image using a 4 µm^2^ minimal threshold, as used in a similar analysis (Ma et al., 2020). The results obtained were screened manually to correct for any false-positive detection and other segmentation errors.

### FIB-SEM sample preparation

*Drosophila* 3^rd^ instar larvae were dissected in SIM. Each pair of dissected glands was transferred to an aluminum planchette (0.15 mm/0.15 mm cavity, 3 mm dia.; Wohlwend GmbH) pre-coated with 1-Hexadecane and submerged in freezing solution (either a mixture of 10% FBS/10% BSA or 20% Ficoll PM70 (F2878-50G; Sigma) in SIM). The planchette was placed on a Whatman No.1 filter paper before it was covered with another 1-Hexadecane-coated planchette (0.3 mm/0.0 mm cavity, 3 mm dia.; Wohlwend GmbH) flat side towards the sample. The closed sample was quickly transferred to the sample holder middle plate of a high-pressure freezer (EM-ICE; Leica Microsystems) and frozen. Frozen samples are stored in liquid nitrogen until an automatic freeze substitution (AFS) and resin embedding procedure. For AFS, the samples were processed in an AFS-2 machine with an FSP pipetting robot (Leica Microsystems according to the protocol in (Ronchi et al., 2021). Briefly, samples were incubated in a freeze-substitution solution (FS; 0.1% uranyl acetate in anhydrous acetone) at -90 °C for 72 h before a gradual (3 °C/h) increase of temperature to -45 °C and kept at this temperature for 5h. FS solution was exchanged with anhydrous acetone with 3 washes. Resin embedding was performed through a gradual increase of Lowicryl HM20 (EMS; #14340) percentage: 10%, 25%, 50%, 75%, and 100% of Lowicryl HM20 in acetone. During 50% and 75% Lowicryl mixing, the temperature was increased gradually (1.67°C/h) to -35 °C and -25 °C, respectively. 100% Lowicryl HM20 solution was exchanged 3 times each exchange for 10h incubation at -25 °C. UV-mediated polymerization was performed at -25 °C for 48 h before temperature was increased to 20 °C (5 °C/h) while the UV lamp was on. The embedded samples are then covered by foil in a chemical hood to cure until their color changes from pink to transparent.

### Block-face confocal imaging

Before imaging the block using confocal microscopy, the block surface was trimmed using a steel blade, keeping the embedded tissue only. The surface of the block was polished with a wet diamond knife (DU3530, Diatome), and sections were stained with toluidine blue and screened on an Olympus CKX53 light microscope using 4x 0.13 NA and 20x 0.4 NA air objectives to verify that the tissue is exposed close to the lumen area. Blocks were then mounted with their surface facing down on a small drop of DPBS inside a 10 mm glass bottom 35 mm imaging plate. The block was pressed flat on the glass and secured using Plasteline. Using the same Olympus IX83 spinning disk confocal microscope (see Confocal imaging section), blocks overview in brightfield was acquired using a 4x 0.16 NA air objective before blocks were mapped using confocal imaging with 20x 0.75 NA air and 60x 1.4 NA oil immersion objectives.

### FIB-SEM block preparation

The block volume was reduced using a fine saw and sanding paper before mounting on an aluminum specimen mount for SEM (EMS; #75220) using double-sided carbon tape (EMS; #77816). The block was then covered with additional carbon tape from the edge of the surface to the specimen mount. Additionally, 2-4 straps of copper conductive tape (EMS; #77802) were placed on top of the carbon tape. The sides of the block were covered with colloidal silver paste (EMS; #12630) and left to dry overnight at RT. One day before imaging, the specimen was coated with a 10nm-thick layer of Iridium using a compact coating unit (Safematic; CCU-010). Iridium coating was performed under 8e-3 mbar process pressure with an inert environment (Ar) and 35 mA. The specimen was placed overnight under the chamber vacuum in a Crossbeam550 FIB-SEM (Carl Zeiss).

### FIB-SEM tomography

All FIB-SEM tomography data was collected using Crossbeam550 or Crossbeam540 machines (Carl Zeiss), and all procedures were performed with SmartFIB or Atlas3D 5 (Carl Zeiss). ROIs were located using SEM at 5-15 kV, 350 pA. The stage was then tilted to 54° to face the ion beam stage height, adjusted to SEM and FIB coincidence point (5mm working distance). An ion beam-induced platinum deposition was performed using a 30 kV, 0.7-1.5 nA probe. The trench is milled using 30 kV, 15 nA, and the cross-section was polished with a 30kV, 3nA probe. Slices of 10nm-thickness were removed using a 30 kV, 0.7-1.5 nA ion probe, and the cross-section was imaged using an electron beam at either 2 kV, 0.35 nA, or 1.5 kV, 1 nA. In stacks acquired using SmartFIB, the signal was collected using either a secondary electrons type two detector (SE2), an energy-selective backscatter detector (EsB), or any mixed signal between them. In stacks acquired using Atlas3D 5, the signal was collected using EsB. Pixel dwell time varied between 4µs and 8µs, and the voxel size ranged between 10 nm x 10 nm x 10 nm and 15 nm x 15 nm x 15 nm, depending on the size of the acquired volume.

### FIB-SEM tomography stack processing

FIB-SEM stacks were aligned via Scale-invariant feature transform (SIFT) using linear stack alignment with the SIFT plugin for Fiji (Lowe, 2004; Schindelin et al., 2012), allowing only translation of images with respect to one another. The aligned stacks were cropped to remove the excess background. Manual cropping of horizontal lines in Fourier space was done to remove vertical strips caused by the ion beam on the cross-section. CLAHE was used in Fiji built-in function local contrast enhancement to increase contrast if needed (Schindelin et al., 2012; Zuiderveld, 1994). Binning of 2- fold or 4-fold was used to reduce the data size for easier processing during correlation and 3D visualization procedures.

### Pseudopodium dimension quantification

The pseudopodium dimensions were measured in Fiji (Schindelin et al., 2012) and saved using the ROI manager tool. Only LSVs acquired entirely in the FIB-SEM stack were used to obtain pseudopodium dimensions. The middle plane of the pseudopodium was found by parsing through the z-axis, and a line was measured at the base of the pseudopodium. The width at the base of the pseudopodium was defined at the point of curvature change from positive to negative on both ends. A straight, perpendicular line from the width line to the tip of the pseudopod was measured to measure the height. The ratio between the height and width of each pseudopodium was calculated by dividing the height by the width. This was made to measure the pseudopodium proportion and cancel out the population variability effects. A spherical cup calculation was made to estimate the surface area of each pseudopodium, assuming that a pseudopodium is a centrosymmetric object.

### Polarity analysis

Segmented data was used to extract the profile of each segmented vesicle (segmented was done using Amira, Thermofisher Scientific). For each segmented vesicle, an ellipsoid fitting was made using Fiji (Schindelin et al., 2012). The segmented data and physical parameters from the ellipsoid fitting were used as input for a script written in C#. The script generated the outline of each segmented LSV and an ellipsoid fitting. The RGB values of each segmented LSV were used as a unique identifier by the script and were numbered. Then, a minimal pixel difference between the LSV profile and ellipsoid, and the minimal angle between the pseudopodium vector and a dent vector were inserted into the graphic user interface. The script then compared the difference between the points on the LSV profile and the ellipsoid fitting and generated a vector from the center of the LSV to the point with the largest distance between the profile and the ellipsoid, for both pseudopodia and dents in the neighboring LSV. The deviating angle was defined to validate the dent in the neighboring LSV as a tool to reduce false detection. The output from the analysis was a table with all pseudopodia and dents detected and the angle in clockwise coordinates (12 o’clock = 0°, 3 o’clock = 90°, etc.). The angles were then converted to conventional polar angle coordinates, and only vectors of pseudopodia were extracted for plotting. Manual validation of the vector angle was calculated in Fiji (Schindelin et al., 2012) by using the line tool and extracting the angles using the built-in measure function. Additionally, a manual count of all undetected pseudopodia was conducted to determine the percentage of false negatives in the analysis.

The direction of the lumen was estimated by the range of angles with respect to the center of an image. In the volume used for the polarity analysis the lumen was stretched between 240°-270° (polar angle coordinates). This area was highlighted in gray on the plot of the average vector angle to indicate the range of angles in which the lumen surface stretches (Fig. S1B-C).

### Correlation of confocal microscopy and FIB-SEM data

Before correlation, confocal stacks were processed in Fiji, in case of low signal-to-noise ratio, by application of background subtraction. Briefly, two median filters were applied on two copies of the stack (2 pixels and 50 pixels median filters), and the resulting filters were subtracted from one from the other using the image calculator in Fiji. Data correlation was performed using either eC-CLEM v2 plugin for Icy (De Chaumont et al., 2012; Paul-Gilloteaux et al., 2017; Potier et al., 2021) or by Bigwarp plugin for Fiji (Bogovic et al., 2016). In both cases, the image transformation algorithm was picked based on the most restrictive transformation that provided a correlation that properly overlayed the fiducial markers in both volumes. In all correlations in the *Drosophila* larval salivary gland, LSVs were used as fiducial markers for correlation. Either similarity or affine transformation was used. The correlation was based on 15-30 fiducial markers, sparsely spread over the entire volume, unambiguously identified in the confocal stack and the resliced FIB-SEM stack with the same orientation as the confocal stack. Correlation was refined by providing more fiducial markers and removing those that contributed to a larger error in the transformation. In cases where correlation was performed using eC-CLEM v2, a predicted error map of the entire volume was calculated to give a mathematical indication for correlation precision (Potier et al., 2021; Scher et al., 2022).

### Sample preparation and pull-down assay

Four batches, each of 30 pairs of salivary glands were dissected out of 3rd instar larvae of UAS-MIM-Emerald under C135-GAL4 expression or yellow white (yw) in SIM. Glands were gently washed with PBS before being incubated in crosslinking solution (1 mM dithiobis(succinimidyl-propionate); (DSP) in PBS). Following 40 min of incubation at RT, Tris-Cl pH 7.5 was added to a final concentration of 100 mM to quench the crosslinking. The glands were then transferred to a clean 2 mL Dounce homogenizer in 250 µL radioimmunoprecipitation assay buffer (RIPA; 150 mM NaCl, 50 mM Tris-Cl pH 7.5, 1 mM EDTA pH 8, 1% sodium deoxycholate, 0.1% SDS, 1% NP-40), freshly supplemented with Halt™ Protease inhibitor cocktail (Thermo Fisher Scientific). Homogenization was performed by 20 Dounce strokes on ice, followed by transferring the lysate to a clean 1.5 mL tube and incubation for 1 h on ice before clearing by centrifugation at 10,000xg for 10 min in 4°C. The supernatant was then collected into clean tubes and incubated with 50 µL of magnetic microbeads conjugated to α-GFP antibody (µMACS™ GFP isolation kit; Miltenyi Biotech) for 1.5 h in 4°C. Columns were placed in a Multi™MACS M96 separator (Miltenyi Biotech), equilibrated in lysis buffer, then pull-down samples were then loaded onto columns and let flow. Columns were washed 3 times in 800 µL of wash buffer #1 (150 mM NaCl, 50 mM Tris-Cl pH 7.5, 1 mM EDTA pH 8, 0.1% sodium deoxycholate, 0.01% SDS, 0.1% NP) followed by 2 washes with wash buffer #2 (150 mM NaCl, 50 mM Tris-Cl pH 7.5, 1 mM EDTA pH 8). Elution of samples from column was achieved through on-column trypsinization by 30 min incubation in 25 μL Elution buffer I (2 M Urea, 50 mM Tris-Cl pH7.5, 1 mM DTT, 5 mg/ml Trypsin), followed by the addition of 100 μL Elution buffer II (2 M Urea, 50 mM Tris-Cl pH 7.5, 5 mM Chloroacetamide). Eluate was collected in a new tube and allowed continued trypsin digestion overnight.

### LC-MS/MS sample processing

Digested and acidified peptides were cleaned up using C18 StageTips with three layers of Empore octadecyl C18, 47 mm extraction disks. LC-MS analyses were conducted using a Dionex UltiMate RSLC 3000 nano-HPLC system coupled with an Orbitrap Exploris 480 mass spectrometer (Thermo Fisher Scientific), employing high-field asymmetric-waveform ion-mobility spectrometry (FAIMS) at a compensation voltage of -45 V in data-independent acquisition (DIA) mode. Peptides were loaded onto an Acclaim PepMap 100 µm × 2 cm C18 precolumn (Thermo Fisher Scientific) and separated on an Aurora Frontier 75 µm ID × 60 cm, 1.7 µm C18 column (Ionopticks) maintained at 50°C. Full MS1 scans were acquired over 400–1200 m/z at a resolution of 30,000 with an AGC target of 300%; DIA fragmentation scans were performed over 450–800 m/z using 38 isolation windows of 9 Th with 1 Th overlap, at a resolution of 45,000 with an AGC target of 3000%. Fragmentation was achieved by normalized high-energy collisional dissociation (HCD) at collision energies of 22%, 26%, and 30%. Peptide separation employed a 110-minute gradient at 0.250 µL/min: 1-43% solvent B (80% [v/v] acetonitrile, 0.1% [v/v] formic acid) over 82 minutes, a 10-minute wash to 98% B, and an 18-minute re-equilibration at 1% B. Raw MS files were converted to .d format and analyzed using DIA-NN (version 1.8.1) with an *in silico* spectral library generated from the canonical *Drosophila melanogaster* proteome (UniProt release 2018_03, taxon ID 7227; 22,006 protein sequences, 14,373 genes) and predicted peptide retention times. The DIA-NN search was configured with precursor and fragment mass ranges of 300–1800 m/z and 200–1800 m/z, respectively, a scan window radius of 5, precursor charge states +2 to +4, and peptide lengths of 7-30 amino acids. Interfering precursor peaks were removed, robust LC (high-accuracy) quantification was applied, precursors were filtered at a q-value threshold of 0.01, and protein grouping was performed at the canonical gene level using a double-pass neural network classifier with per-run mass accuracy optimization and a match-between-runs (MBR) strategy. Downstream statistical analysis was performed in Perseus (version 2.0.11; (Tyanova et al., 2016)). Protein intensities were Log2-transformed and filtered to retain proteins with a minimum of four valid values in the MIM DSP pulldown group. Missing values in the NO MIM DSP control group were then imputed from a normal distribution (width = 0.3, down-shift = 1.8 standard deviations, applied separately per column). Protein isoforms were collapsed to the gene level by taking the median intensity across isoforms. Differential abundance was assessed using a one-sided two-sample Student’s t-test testing for enrichment in the MIM DSP pulldown over the NO MIM DSP control, with permutation-based FDR control (FDR < 0.01, S0 = 0.250 randomizations).

### GO terms analysis

The resulting protein IDs list from the MS analysis of the DSP dataset were used to extract GO terms using a Python 3.12 script in PyCharm community edition (JetBrains). The script used pandas, scipy and bioPython libraries and was written with the assistance of ChatGPT based on the GPT-5.3-mini model (OpenAI). First, a list of all enriched proteins IDs in the DSP dataset were fed to the script. GO terms for each protein ID were extracted using the UniProt database. Then, the script counted the number of instances each GO term appeared, and the script generated an Excel file with the list of all protein IDs and their GO terms and another sheet with the GO terms count. The top GO terms counts that appeared in the enriched dataset were included in the plot. Based on the GO terms, we conducted an Over-Representation Analysis (ORA) between the GO terms count of the 746 significantly enriched proteins and the entire dataset of proteins identified by the mass-spectrometry-based analysis (4573 proteins). Based on the ORA, a dot plot was generated using matplotlib library to represent the 30 GO terms values with the lowest false-discovery rate. The size of the dots represents the number of proteins among the enriched proteins that are included in the GO term.

### Immunofluorescence

We followed the procedure described by Boda et al. (Boda et al., 2023). Briefly, the glands were dissected and permeabilized in 0.1% Triton X-100 and 0.5% sodium deoxycholate in DPBS without calcium and magnesium (DPBS wo/wo) for 40 seconds at RT. Glands were then fixed using 4% PFA in DPBS wo/wo for 30 min followed by two washes with DPBS wo/wo of 15 min at RT. Then, the samples were re-permeabilized in 0.3% Triton X-100 in PBS for 10 min at RT. Blocking was done with 10% BSA and 5% normal goat serum (NGS) in PBT (DPBS supplemented with 0.1% Tween x-100) for 30 min at RT. The samples are then incubated with the primary antibodies mouse α-V5 1:200 (R960- 25, Thermo Fisher Scientific), mouse α-Flag 1:00, and Phalloidin-Atto-640 1:2000 (AD 641-81; ATTO-TEC) in 10% BSA 5% normal goat serum (NGS) in PBT for 2 days at 4°C and then washed two times for 15 min in PBT. Before secondary antibody incubation, the samples were blocked for 30 min in 10% BSA in PBT at RT, followed by the secondary antibody incubation Alexa Fluor 555 goat α-mouse 1:1000 (A-21424; Thermo Fisher Scientific) incubation for 4h at RT. Samples are then washed and mounted with Fluoromount (F4680, Merck) on 35 mm imaging plates with a 10 mm glass bottom well.

### Graphs and statistics

The *Drosophila* growing conditions and experiment procedure were similar in the different experiments. Each experiment included at least 3 different individuals. The detailed analysis for each quantification is referred to in the individual sections of the materials and methods. All plots and statistical tests in this work were done using GraphPad Prism version 10 for Windows. P values: ns > 0.5, * < 0.1, ** < 0.01, *** < 0.001, **** < 0.0001. The specific details on sampling size and the statistical test performed are listed in the relevant figure legend.

### Figures and illustrations preparation

All figures were made in Adobe Illustrator version 30.3 (64-bit) for Mac. Some of the illustrations use assets from Biorender (Biorender.com)

## Supplemental materials

Sup. Figures S1-S6

Movies: Movies1-5

Table 1: Mass spectrometry-based proteomic results.

Table 2: Fly lines.

## Data Availability

Mass spectrometry raw data and DIA-NN output files (protein group and precursor matrices) have been deposited in the PRIDE database (ProteomeXchange consortium) and will be publicly available upon acceptance; the accession number will be provided upon receipt.

## Acknowledgments

This work is dedicated to the memory of our dear friend and colleague Ben-Zion (Benny) Shilo (1951-2025). Benny’s scientific insight and boundless curiosity were integral to this study and to the environment in which it grew. We deeply miss his wisdom, warmth and joy in discovery, and regret that he did not live to see this work published. We thank all members of the Avinoam Lab for fruitful discussions. We also thank Prof. Jullie Brill for the Sgs3-DsRed fly line and the UAS-Tsp29Ta line, Prof. Eli Arama for the UASp-CD63-GFP and UASp-CD63-RFP, Dr. Helen Zimmer for the MIM^null^. FIB-SEM data acquisition was conducted at the Irving and Cherna Moskowitz Center for Nano and Bio-Nano Imaging at the Weizmann Institute of Science and at the electron microscopy core facility (EMCF) at the European Molecular Biology Labs. We thank Mr. Moshe Varsano for writing the polarity analysis script. O.A. is an incumbent of the Miriam Berman presidential development chair

---

**Supplementary figure 1:**
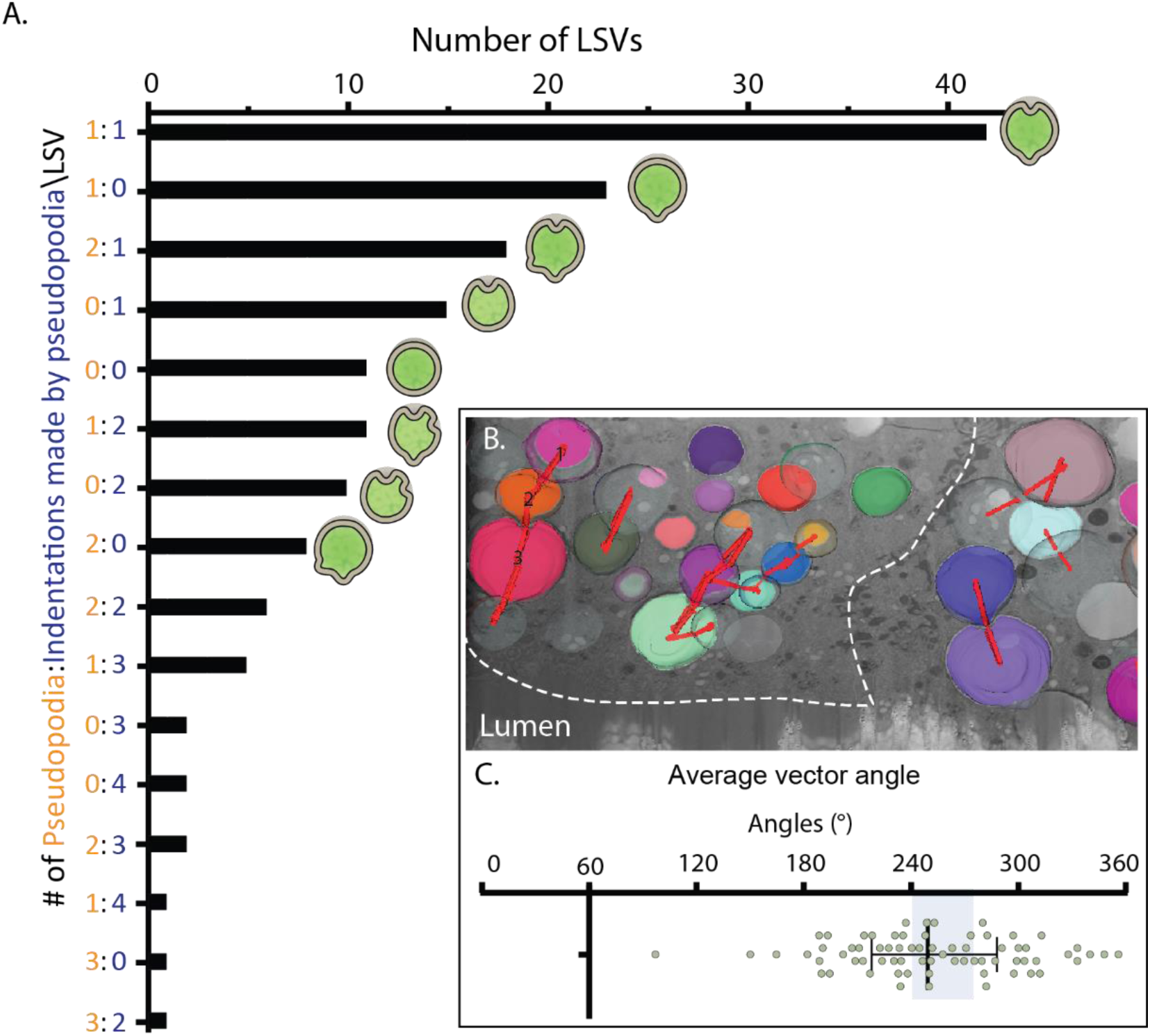
**A.** Histogram of the quantified LSVs from Fig. 1F, showing the distribution of LSVs with different amounts of pseudopodia and dents formed by pseudopodia. As indicated, most LSVs have one pseudopodium projected from them and one dent. Only seldomly were there LSVs with more than 2 pseudopodia and 2 dents. **B.**-**C.** Analysis of the polarity in control salivary glands. We analyzed one FIB-SEM volume to assess the polarity of the pseudopodia (see materials and methods section). **B.** LSVs are numbered and color-coded and outline of the LSV and an ellipsoid is fitted into every LSV. The image shows a volume rendering of the LSVs on top of the cross-section. Red lines connected between the LSVs indicate the vectors from the center of an LSV to the tip of the pseudopodium it projects. From the vectors, we calculated the vector angles. **C.** The angles extracted from vectors. Pseudopodia vectors are facing the direction of the apical membrane (gray block behind the plot).

**Supplementary figure 2:**
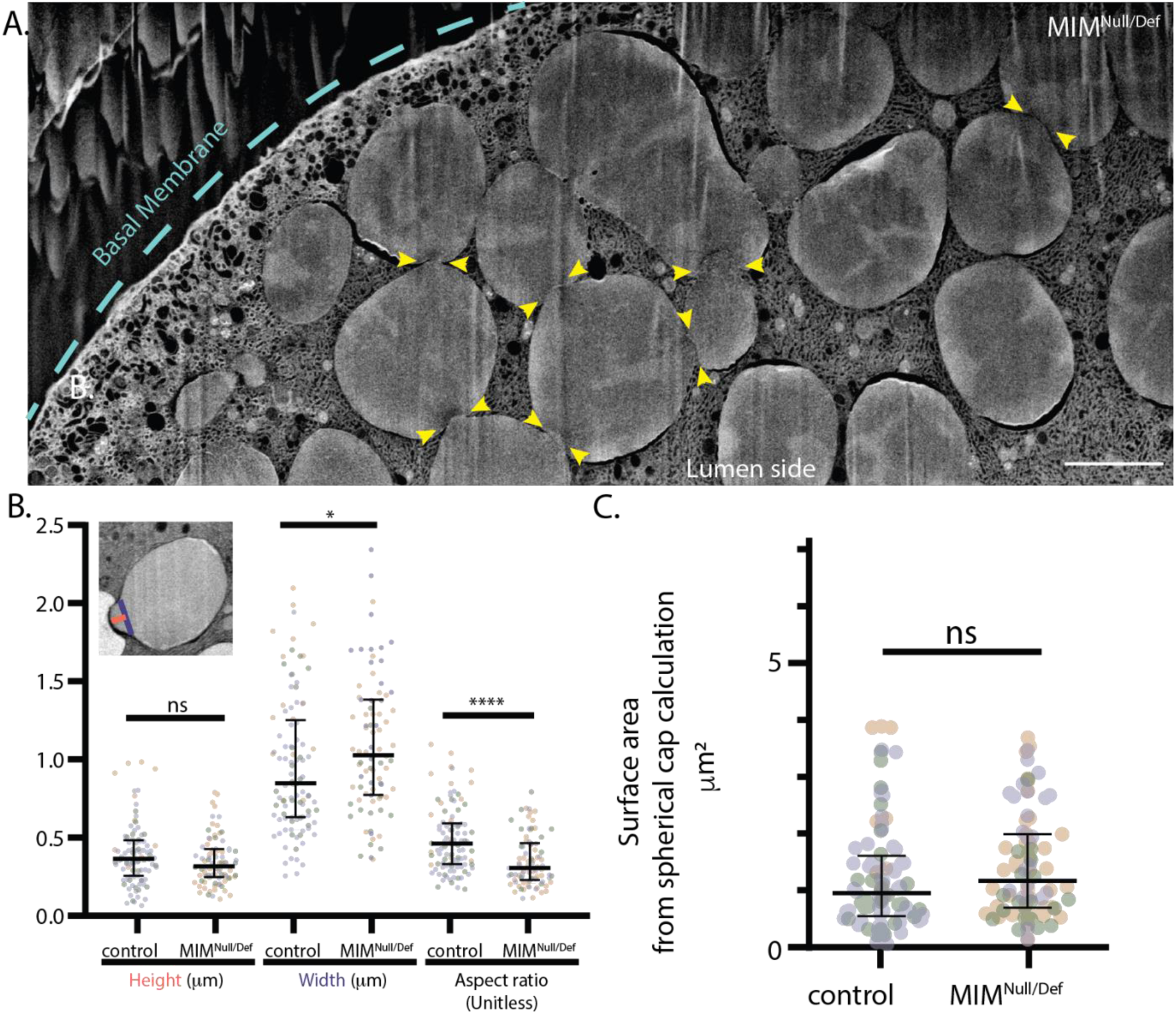
**A.** A representative FIB-SEM tomography slice through a secretory cell from a MIM^Null/Def^ mutant salivary gland showing vesicular pseudopodia (yellow arrowheads). The pseudopodia appear to face towards the basal rather than the apical membrane and their shape is altered slightly. Scale bar: 5 μm. **B.** Quantification of the height, width and aspect ratio of vesicular pseudopodia in in glands expressing Sgs3-GFP and LifeAct-Ruby, derived from control and MIM^Null/Def^ mutant larvae. While the difference in height is not significant, the width of vesicular pseudopodia at the base is larger in MIM-null^Null/Def^ compared to the control. The difference in pseudopodia morphology increases when comparing the aspect ratio between the MIM^Null/Def^ mutants and the control (control: *N*_*larva*_=3, *n*_pseudopodia_=101, MIM^Null/Def^: *N*_*larva*_=3, *n_pseudopodia_*=84). The aspect ratio takes into account the variability in the population, and depicts the change of pseudopodia themselves. These results suggest that MIM is involved directly or indirectly in supporting vesicular pseudopodia morphology. **C.** Pseudopodia surface area comparison between control and MIM^null/Def^ salivary glands expressing Sgs3-GFP and LifeAct-Ruby show no significant difference in the spherical cup calculation, which suggests that the amount of membrane in pseudopodia of both genotypes is similar despite the difference in pseudopodia aspect ratios.

**Supplementary figure 3:**
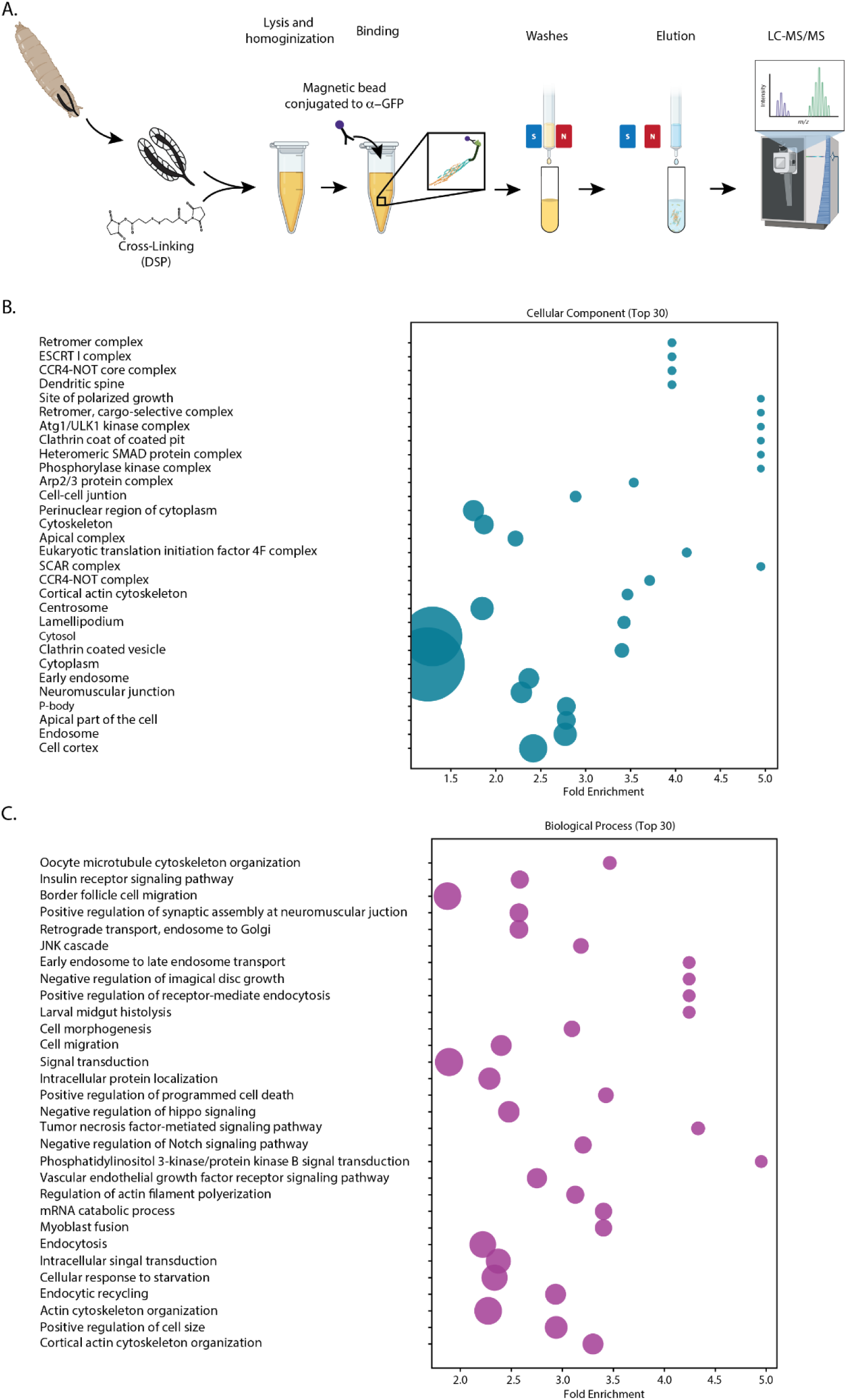
**A.** Schematic representation of the pull-down workflow. Glands over-expressing MIM-Emerald under UAS control were dissected in Schneider’s Insect Media. After dissection, the salivary glands are cross-linked using the permeable crosslinker Dithiobis(succinimidylpropionate) (DSP) before lysis and homogenization. The lysate is then incubated with α-GFP conjugated to magnetic microbeads and loaded onto a column. The sample is then put in a magnetic field to hold the beads, which are then washed to remove non-specific adsorbed proteins, before elution. The eluate is processed and analyzed using LC-MS/MS. **B.-C.** Gene Ontology (GO) enrichment analysis of the significantly enriched proteins (746 proteins) over the background of all proteins identified in the mass-spectrometry based analysis (4573 proteins) of the MIM-Emerald pull-down. Dot plots showing the top 30 enriched GO terms based on an Over-representation analysis (ORA) for **B.** Cellular components and **C.** Biological processes. The dot size corresponds to the proportion of identified proteins associated with a term. The 30 GO terms presented were scored with the lowest false-discovery rate (FDR). The fold-enrichment indicates that the term appears more frequently within the proteins enriched in the MIM-Emerald pull-down than would be expected by chance. (Figure elements adopted from BioRender: Biorender.com).

**Supplementary figure 4.**
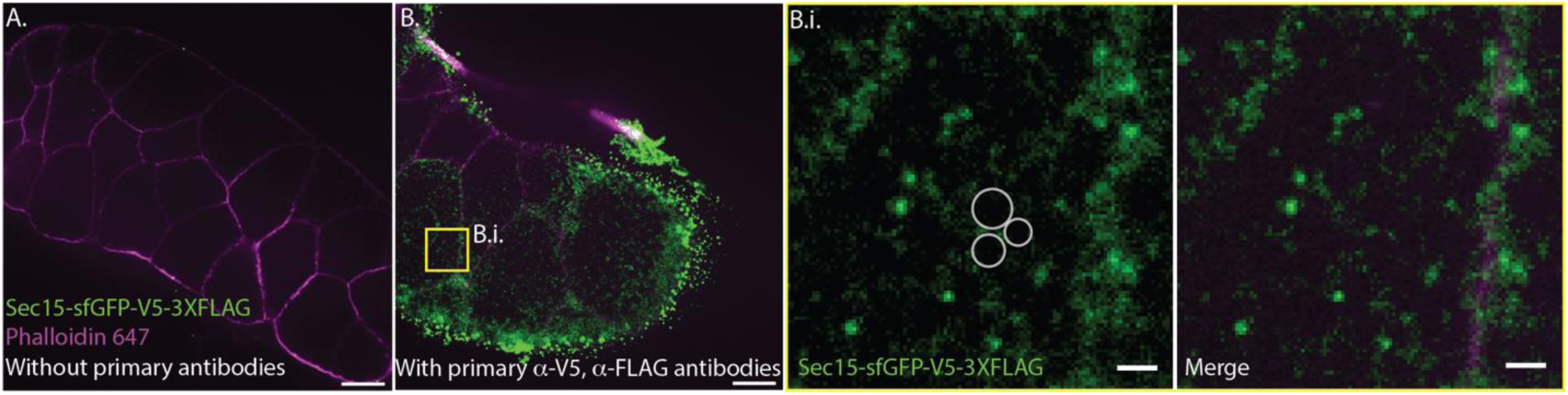
Sec15 localization between the LSVs is apparent at endogenous expression levels. **A.** Control immunofluorescence assay without primary antibodies for the V5 and 3XFLAG tags showing no signal in green. Actin is stained with phalloidin (magenta) for additional context of the tissue. **B.** Using primary antibodies against V5 and 3XFLAG yield signal, indicating Sec15 localization. The inset **B.i.** is an enlarged image of the yellow box, showing Sec15 localizing in between the LSVs (depicted by three circles), even at endogenous expression levels. Scale bars **A.-B.** 20μm, **B.i.** 5 μm.

**Supplementary figure 5.**
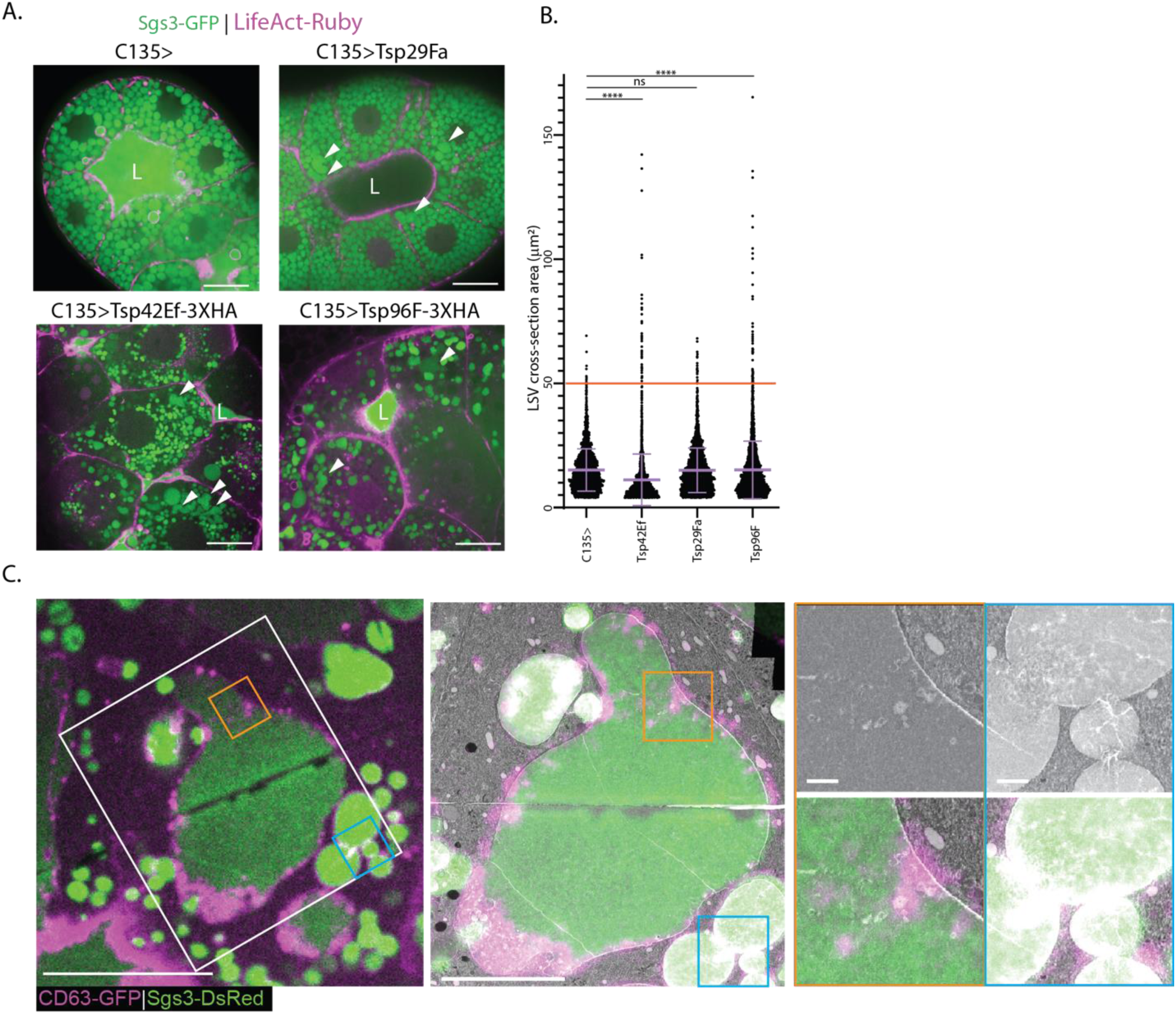
Over-expression of TSPs under the C135-GAL4 driver causes deformations in LSVs morphology to different severities. **A.** Representative images of over-expressed TSPs in salivary glands expressing Sgs3-GFP and UAS-LifeAct-Ruby. All over-expressed TSPs exhibit large deformed LSVs (white arrowheads). Scale bar: 20 µm. **B.** Quantification of the LSV cross section area of the over-expressed TSPs from **A**. LSV cross-section area of the over-expressed TSPs show an increase in the number of LSVs with a cross-sectional area larger than 50 µm^2^ (red threshold line), particularly in Tsp42Ef and Tsp96F. Each over-expressed TSP exhibit different penetrance, where C135>UASTsp42Ef3xHA caused phenotype in all observed glands (n=9) while C135>UAS-Tsp29Fa and C135>UADTsp96F3xHA both exhibit 56% and 50% penetrance (n=9, n=6, respectively). Because the oversized LSVs are the result of fusion between smaller vesicles, calculating the number of average-sized vesicles that make the oversized vesicles suggested 7% (185 vesicles) and 8% (308 vesicles) increase in number of vesicles in Tsp42Ef and Tsp96F over-expression, respectively. Number of different salivary glands of each phenotype, N=3, number of vesicles, n=2462, 2605, 2861, 3867 (C135>, Tsp42Ef, Tsp29Fa, Tsp96F, respectively). Two-tailed non-parametric t-test using the Kolmogorov-Smirnov method. **C. Left**: Block-face FM depicting a cell of interest and an oversized LSV filling most of its volume, and the area acquired using FIB-SEM (white rectangle). **Middle**: 3D-CLEM Overlay of transformed confocal micrographs depicted to the left and FIB-SEM stack of a secretory cell with large LSV. FIB-SEM image resliced to present the XZ plane from the FIB-SEM (which matches the XY in confocal). The CD63-GFP signal corresponds to membranal structures inside the lumen of the LSV (orange) but does not appear to be enriched in vesicular pseudopodia (teal). Scale bars: 20 µm, insets 1 µm.

**Supplementary figure 6.**
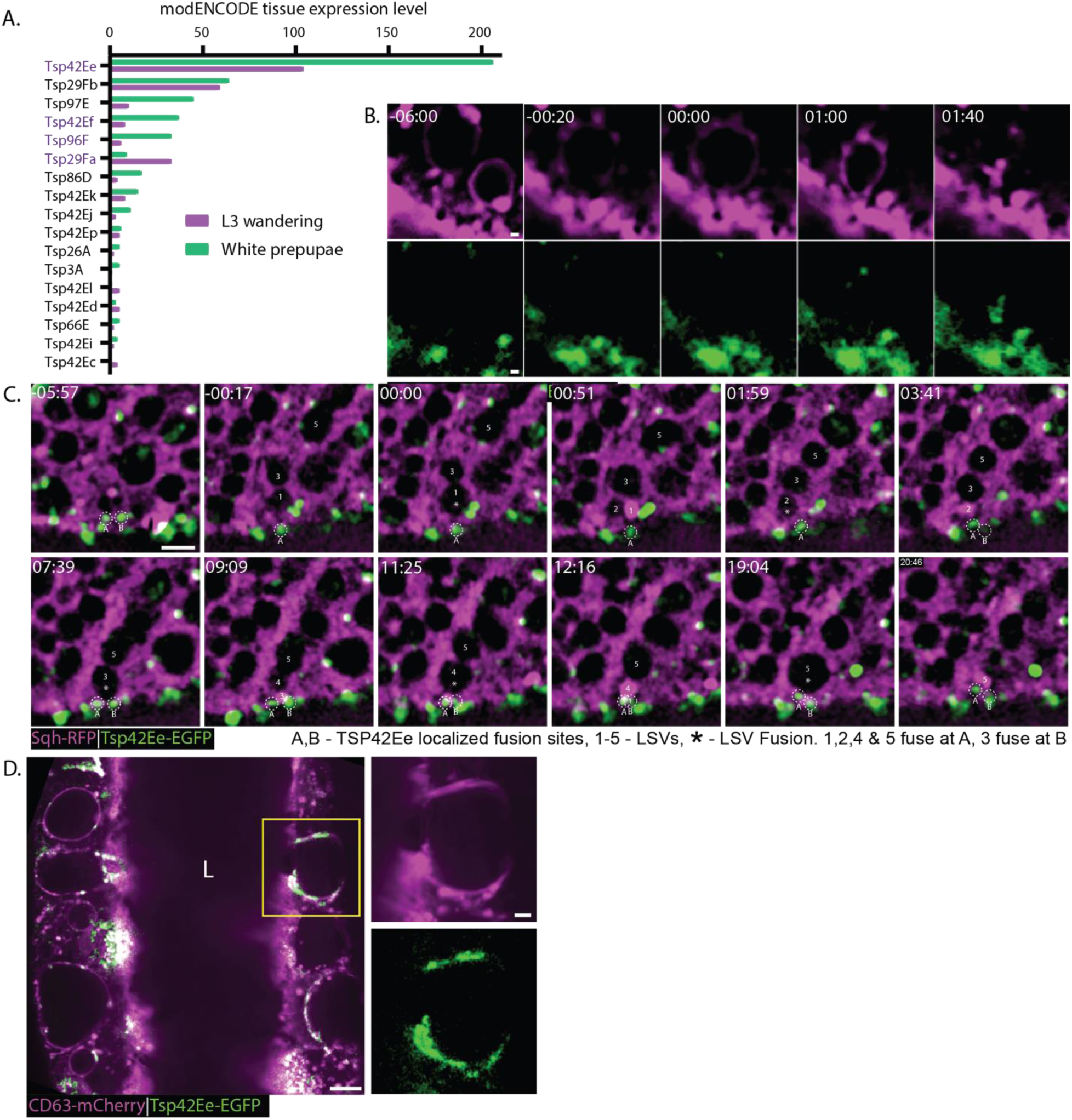
**A.** Expression levels of tetraspanins in the salivary glands of third instar larval stage (green) and the white prepupae stage taken (purple) from The Flybase. Tsp42Ee exhibits the highest expression level in both stages. **B. Corresponds to merged image in** Fig. 6A. Time-lapse series (presented as confocal intensity-projections), of representative LSVs from salivary glands expressing both the MIM-mScarlet (Magenta; under UAS control) and the Tsp42Ee-eGFP (Green; under endogenous promotor; BDSC_51588) markers. **C.** An example of a time-lapse series from salivary glands Sqh-mCherry (magenta) and Tsp42Ee-EGFP (green) under their endogenous promoters. LSVs can be seen fusing to two close domains on the apical surface (marked a and b) where Tsp42Ee is localized. LSVs are numbered 1-5, fusion events are marked with an asterisk. Time mm:ss; Scale bar: 5 µm. **D.** Representative image from confocal slices projected in z of salivary gland expressing CD63-mCherry (magenta; overexpression using UAS driven by C135-GAL4) and Tsp42Ee-EGFP (green; endogenous expression). The overexpression of CD63 causes a severe compound exocytosis phenotype, in which Tsp42Ee is largely absent from the apical membrane and appears on the large vesicles which are the result of the compound exocytosis. Scale bars: 20μm, inset: 5 μm.

## Supplementary movies 1-5

### Supplementary Movie 1

The segmented FIB-SEM stack shown in Figure 1, including the vector analysis described in Supplementary Figure 1.

### Supplementary Movie 2

Time-lapse series of an LSV fusing with the apical surface in salivary glands expressing Sgs3-DsRed (green) and GFP-Sec15 (magenta; under UAS control). Sec15 puncta are present during secretion at the interface between the LSV and the apical membrane.

### Supplementary Movie 3

Time-lapse series of representative LSVs from salivary glands expressing both the MIM-mScarlet (Magenta; under UAS control) and the Tsp42Ee-EGFP (Green; under endogenous promotor; BDSC_51588) markers. Fusion - white asterisk (Vesicle swelling), LSV - White dashed transparent polygon line (marked using the MIM-mScarlet channel), MIM puncta at fusion site (LSV)- Yellow dashed circle, Tsp42Ee puncta at fusion site (apical) - Blue dashed circle. Time mm:ss; relative to fusion. Scale bar: 1 µm.

### Supplementary Movie 4

Tsp42Ee localized to distinct fusion sites on the apical membrane. Salivary gland expressing Sqh-mCherry (magenta) and Tsp42Ee-EGFP (green) under their endogenous promoters. LSVs can be seen by the shaded areas in Sqh-mCherry signal. Time-lapse series of a small area from a secreting gland. showing that four vesicles fuse to the same area on the apical membrane (fusing vesicles marked in asterisk) within less than 20 minutes.

### Supplementary Movie 5

Another time lapse series of the salivary gland expressing Sqh-mCherry (magenta) and Tsp42Ee-EGFP (green) under their endogenous promoter showing consecutive fusion events to Tsp42Ee-positive membrane domains.

