## Supplementary Table 1 for "Vesicular pseudopodia define the fusion site on large secretory vesicles of the *Drosophila* salivary glands"

**Supplementary Table 1:** **Proteins enriched in a MIM-Emerald pull-down assay**. Mass-spectrometry based proteomics results from an analysis of immunoprecipitates using an anti-GFP antibody from *Drosophila* salivary gland lysates. Significant potential interactors were identified using a one-sided Student's t-test (FDR <0.01, S0=0.25).

| Gene name | Protein IDs | Student's T-test Significant MIM_NO MIM | -Log10(p-value) | Student's T-test q-value MIM_NO MIM | Student's T-test Difference MIM_NO MIM |
| --- | --- | --- | --- | --- | --- |
| mim | A0A0B4JD47;A0A0B4K7T1;A0A0B4KED3;A1Z6S0 | + | 6.813763379 | 0 | 10.31506252 |
| CG14261 | A0A0B4LHR8;Q8IMP2 | + | 4.977579587 | 0 | 7.696367979 |
| Alms1a | Q5BI31 | + | 6.415303194 | 0 | 6.460788727 |
| w | P10090 | + | 4.74015556 | 0 | 5.396086454 |
| Dmel\CG3397 | Q9VGF1 | + | 3.3920699 | 0.001361345 | 4.206862688 |
| Dmel\CG12661 | Q9W3A7 | + | 5.123518273 | 0 | 4.071716785 |
| Dmel\CG8087 | Q9VFI5 | + | 4.528045253 | 0 | 4.060343981 |
| lva | Q8MSS1 | + | 3.844060743 | 0.000676056 | 3.902343273 |
| Abi | A0A0B4K774;A0A0B4KH51;Q9Y0S9 | + | 4.036361123 | 0.000521739 | 3.809606552 |
| Hip1 | Q8MQJ8 | + | 2.386199922 | 0.008610951 | 3.807483196 |
| unc-13 | Q9V483;X2J9E1 | + | 3.150724627 | 0.002139241 | 3.804495573 |
| Cyfip | Q9VF87 | + | 3.522485071 | 0.001327103 | 3.788024664 |
| CG10857;CG12604 | Q7YU16;Q9VZM9 | + | 4.503127537 | 0 | 3.783102989 |
| GstE9 | Q7K8X7 | + | 5.173590992 | 0 | 3.780598164 |
| SCAR | F2FB81;H5V8D0;M9NCT2;M9PCP4;Q9VKM2;X2J5L4 | + | 4.416812798 | 0 | 3.778280497 |
| Dmel\CG11248 | Q9VNZ7 | + | 5.441727564 | 0 | 3.767258167 |
| Dmel\CG9795 | Q59E01;Q8IPM6 | + | 5.092099708 | 0 | 3.677119493 |
| Dmel\CG14756 | Q7JRA5 | + | 3.771390794 | 0.001073171 | 3.647732496 |
| Smyd4-1 | A1Z7W1 | + | 5.12353413 | 0 | 3.605746746 |
| tna | A0A6J3SHB9;Q7JR52;Q7KUE4;Q9VTB8 | + | 2.776129094 | 0.00397541 | 3.597868919 |
| ec | Q1W9P9 | + | 4.412299036 | 0 | 3.55515027 |
| gukh | Q9VE13 | + | 2.846479477 | 0.003649123 | 3.498116255 |
| Cortactin | Q9VDF4 | + | 3.559821345 | 0.001223301 | 3.49729991 |
| Tab2 | A0A0B4KG87;Q6NQZ7 | + | 3.754345536 | 0.001060241 | 3.420736074 |
| Plp | M9ND93;M9NFV3;M9PCJ5;M9PCK2;M9PFA8;M9PFJ2;M9PFP0;M9PI63;Q400N2 | + | 4.868793209 | 0 | 3.417477369 |
| lute | A0A0B4KGU2;C7LAF3;Q9VEL0 | + | 4.537310116 | 0 | 3.392940521 |
| Dmel\CG6511 | Q9VSP0 | + | 6.009300204 | 0 | 3.37010169 |
| sowah | B7Z0B0;B7Z0B1 | + | 2.844609469 | 0.003676149 | 3.347683191 |
| PAN3 | Q95RR8 | + | 5.415032001 | 0 | 3.34732914 |
| GEFmeso | A1ZBA1 | + | 2.459532056 | 0.007255385 | 3.34506011 |
| cno | A0A0B4KF82;Q8IPP3;Q9VN82 | + | 3.274402195 | 0.001677903 | 3.339840651 |
| cher | A0A0B4KGT8;Q9VEN1 | + | 2.751099086 | 0.004160643 | 3.330649853 |
| Nadsyn | Q9VYA0 | + | 3.251570602 | 0.001833935 | 3.275883198 |
| ine | Q9VR07 | + | 2.849200638 | 0.003673289 | 3.264708042 |
| NKAIN | A6MHQ4 | + | 2.602717148 | 0.005188049 | 3.260283947 |
| homer | O96607;Q0E8S7 | + | 2.343923039 | 0.009402235 | 3.246723175 |
| Evi5 | Q9VYY9 | + | 4.162686376 | 0.000194175 | 3.230899334 |
| PAN2 | A1Z7K9 | + | 5.896631443 | 0 | 3.219656229 |
| cana | X2J5P0 | + | 3.492610435 | 0.001314815 | 3.20344758 |
| Hem | P55162 | + | 3.39266116 | 0.001367089 | 3.184405327 |
| CG3857 | O46080 | + | 4.614157581 | 0 | 3.174891472 |
| Dif | P98149 | + | 4.785241501 | 0 | 3.141366005 |
| Mlc-c | P54357 | + | 2.682342033 | 0.004604915 | 3.140138626 |
| Mer | Q24564 | + | 2.568167465 | 0.005689949 | 3.136095524 |
| ASPP | A0A0B4KF17;Q86BG1;Q9W2J2 | + | 2.466088029 | 0.007271028 | 3.131387949 |
| Dlg5 | M9PD90;Q9VKG8 | + | 3.992514767 | 0.000633333 | 3.093524694 |
| LP23408p | Q9VAU3 | + | 3.041293566 | 0.002766667 | 3.093425989 |
| Sdb | A1Z9M5 | + | 3.904723586 | 0.000727273 | 3.081284523 |
| GM130 | Q9W289 | + | 2.445447281 | 0.007484067 | 3.061877728 |
| p | Q9VHN9 | + | 4.948594883 | 0 | 3.059018373 |
| Traf6 | Q9W3I9 | + | 2.48222008 | 0.006813187 | 3.056421518 |
| Dmel\CG7029 | A0A0B4KGL8;A0A0B4KH73;A8JR84;Q86B80 | + | 2.508955238 | 0.006165316 | 3.052124023 |
| CCDC53 | Q9VLT8 | + | 2.722141909 | 0.004154739 | 3.049803257 |
| CG5270 | Q9VGP1 | + | 3.144598318 | 0.002155763 | 3.048065901 |
| melt | Q9VS24 | + | 3.317609192 | 0.001401575 | 3.026857853 |
| Golgin104 | Q9VV40 | + | 2.970119616 | 0.002989899 | 3.026806355 |
| Gmap | A0A4D6K5M0;Q9VXU2 | + | 3.358645476 | 0.001370968 | 3.021463156 |
| Dmel\CG18596 | Q9VD22 | + | 4.62325682 | 0 | 3.00902009 |
| Tango1 | Q9VMA7 | + | 3.735611441 | 0.001261905 | 3.005040169 |
| Patronin | A0A0B4KEY4;A0A0B4KFA4;A1ZAU8-3 | + | 3.079728 | 0.002720461 | 2.997813463 |
| Dmel\CG14131 | M9PFB4;Q9VTN4 | + | 3.260881185 | 0.001635036 | 2.996753693 |
| kst | A8JNJ6;M9PBL6;Q7KV70 | + | 2.802923003 | 0.003864693 | 2.967345238 |
| CG3638 | Q9W5A5;Q9W5A5-2 | + | 3.848354978 | 0.000690647 | 2.936988354 |
| Vps16A | Q9VHG1 | + | 3.267134239 | 0.001659259 | 2.936687469 |
| Dmel\CG16719 | Q9VT31 | + | 3.962299825 | 0.000612903 | 2.925079584 |
| Neurl4 | A0A1L4AAD6 | + | 4.575891645 | 0 | 2.920839071 |
| Dmel\CG11307 | B5RJJ4 | + | 3.284188501 | 0.001618321 | 2.913708687 |
| Vps26 | Q9W552 | + | 2.605604536 | 0.005215548 | 2.91023469 |
| Tak1 | Q9V3Q6 | + | 3.608052377 | 0.001081633 | 2.906416655 |
| drk | Q08012 | + | 4.918837352 | 0 | 2.852266788 |
| Abp1 | Q9VU84 | + | 4.488680811 | 0 | 2.832598209 |
| nonC | Q70PP2 | + | 3.627546104 | 0.001115789 | 2.828092575 |
| Dark | Q7KLI1 | + | 2.372489335 | 0.00875461 | 2.824533701 |
| smash | A0A0B4K615;B7Z0T1;E2QCZ8 | + | 3.198408203 | 0.002013245 | 2.821094036 |
| CG46338 | Q7K4V4;Q7K4V4-2 | + | 2.778125083 | 0.00399177 | 2.789261818 |
| bchs | Q9VML2 | + | 2.709245153 | 0.004314779 | 2.774346828 |
| TU37B1 | Q9VJ08 | + | 3.825370092 | 0.000746667 | 2.7724998 |
| Hrs | Q960X8 | + | 2.956812762 | 0.003032419 | 2.763480663 |
| Dmel\CG2556 | Q9VYM2 | + | 2.552062559 | 0.00579798 | 2.74908185 |
| FER;Shark | P18106;P18106-1;Q24145 | + | 2.401246976 | 0.008273256 | 2.732266665 |
| Hsp70A;Hsp70B | P02825;P82910;Q8INI8;Q9BIR7;Q9BIS2;Q9VG58 | + | 3.211621293 | 0.002054054 | 2.729963779 |
| Regnase-1 | Q8T0D9 | + | 2.570909362 | 0.005699659 | 2.72849369 |
| CG7472 | A8JQY9;A8JQZ0 | + | 3.235549216 | 0.002013986 | 2.727599621 |
| qtc | Q9VMU5 | + | 3.424607944 | 0.001333333 | 2.719415426 |
| CG15512 | Q9VAE7 | + | 2.42229457 | 0.007887073 | 2.715867519 |
| Nprl2 | Q9VXA0 | + | 2.873974169 | 0.003356009 | 2.715693951 |
| jvl | A0A0B4K6H8 | + | 4.728055148 | 0 | 2.713861465 |
| AGO2 | Q9VUQ5;Q9VUQ5-2 | + | 3.806380493 | 0.001032258 | 2.710148335 |
| bif | M9PGW8;Q8IR86 | + | 3.105692734 | 0.002531343 | 2.709979057 |
| Prosap | A0A0B4K7Z0;A1Z9K8 | + | 3.393863026 | 0.001372881 | 2.707764387 |
| key | A0A0B4K885;Q9GYV5 | + | 4.695861172 | 0 | 2.700159311 |
| Lrrk | A0A0B4KH16;A0A0B4KHT3;Q9VDJ9 | + | 4.223578896 | 0.0002 | 2.695296526 |
| CG11572;CG11578 | H9XVM5;Q9V4B6 | + | 3.202121721 | 0.002026667 | 2.695104837 |
| Stam | Q9XTL2 | + | 2.756612496 | 0.004136821 | 2.693157911 |
| Exn | Q9VVC6;Q9Y150 | + | 4.332991796 | 0 | 2.691335678 |
| IKKepsilon | Q9V3Y8 | + | 3.146674766 | 0.0021625 | 2.691035032 |
| CG4953 | Q95TN1 | + | 2.739202571 | 0.004126233 | 2.683690071 |
| Dmel\CG3342 | Q9W3Z4 | + | 2.418795578 | 0.007976331 | 2.676846504 |
| Strica | Q7KHK9 | + | 5.479248929 | 0 | 2.674975872 |
| tacc | A0A0B4JD97;A0A0B4K6I7;A0A0B4KFK1;Q8IPN1;Q9VN51 | + | 3.791556136 | 0.001106918 | 2.674208641 |
| Rcd-1 | Q7JVP2 | + | 2.465071314 | 0.007237209 | 2.671869755 |
| brun | Q9VIL0 | + | 3.221807287 | 0.002089347 | 2.665876865 |
| mxt | Q9VR35;Q9VR35-2 | + | 3.874682446 | 0.000711111 | 2.664576769 |
| cnk | Q7KNQ9 | + | 3.263524776 | 0.001653137 | 2.661136866 |
| spdi | Q0KHX7 | + | 2.864293466 | 0.003451685 | 2.655745029 |
| Pepck1 | P20007 | + | 2.34913454 | 0.009307153 | 2.655694962 |
| Ude | Q7KS11 | + | 2.610979354 | 0.005252669 | 2.649732113 |
| GCC88 | Q9VGR1 | + | 2.97047944 | 0.003005076 | 2.647723675 |
| Dap160 | M9ND00;M9PBF6;M9PDC7;Q1WWC9;Q8INU2;Q9VIF7 | + | 3.278981706 | 0.001606061 | 2.647282362 |
| Fhos | E1JI78 | + | 2.993666132 | 0.002934037 | 2.634381771 |
| Dmel\CG2003 | Q9V9S4 | + | 4.110039462 | 0.000566038 | 2.623754263 |
| mask | Q9VCA8 | + | 4.80809255 | 0 | 2.616154432 |
| spn-F | Q9V9Y9 | + | 2.414680697 | 0.008093979 | 2.615826368 |
| egl | Q9W1K4 | + | 4.007597967 | 0.000638655 | 2.613208294 |
| Fak | B7YZL9;E1JGM8;Q0E917 | + | 3.528557087 | 0.001339623 | 2.607723713 |
| eIF4G2 | Q9VCH1 | + | 4.861146062 | 0 | 2.604865551 |
| zip | A0A0B4JD57;A0A0B4JD95;A0A0B4K7Q4;Q59E58;Q59E59;Q99323;Q99323-2 | + | 2.373934316 | 0.008767045 | 2.60171628 |
| Dmel\CG15435 | Q9VR03 | + | 3.060994036 | 0.002813559 | 2.595027208 |
| Dys | Q7YU29;Q9VDW6;Q9VDW6-1;Q9VDW6-2;Q9VDW6-3;Q9VDW6-4 | + | 2.558143051 | 0.005817568 | 2.580343962 |
| RASSF8 | Q9VBP2 | + | 3.286052333 | 0.001624521 | 2.578640461 |
| CG12488;CG42663-RB | A0A0B4KFS6;F0JAQ8 | + | 5.169304666 | 0 | 2.567129374 |
| dnc | P12252;P12252-5;Q9W4T4 | + | 4.050752793 | 0.000535714 | 2.566704035 |
| trio | Q7KVD1;Q8IRI5 | + | 3.237921571 | 0.002028169 | 2.565322399 |
| gw | Q8SY33;Q8SY33-2;Q8SY33-3 | + | 3.223217653 | 0.002111111 | 2.562047958 |
| HPS4 | A1ZAX6;A1ZAX8 | + | 3.431230397 | 0.001345291 | 2.556753635 |
| hebe | Q7K568 | + | 2.581127857 | 0.005543739 | 2.55615139 |
| RhoGAP1A | Q29QE1 | + | 4.09454538 | 0.000555556 | 2.554677963 |
| Slmap | Q9W5R5 | + | 3.5247473 | 0.001333333 | 2.552032709 |
| fliI | Q24020 | + | 2.354854215 | 0.009192686 | 2.547335148 |
| M7BP | A0A0B4K7U0;A1Z718;A1Z720 | + | 3.598028569 | 0.00106 | 2.544079065 |
| Dyrk3 | P83102 | + | 2.596838214 | 0.005293706 | 2.534968615 |
| CG;Dmel\CG2017 | Q86BS2;Q960F7 | + | 4.273299372 | 0 | 2.533614159 |
| siz | Q7KTX2 | + | 5.134813609 | 0 | 2.531242847 |
| DIP2 | M9NDW1;M9PGJ0;Q9W0S9 | + | 3.179851096 | 0.002077419 | 2.530288458 |
| faf | P55824 | + | 2.898885735 | 0.003237209 | 2.525651932 |
| GAPcenA | A8JNN5 | + | 2.670114444 | 0.00472045 | 2.52454257 |
| Dmel\CG34125 | Q0E8T6 | + | 3.838809048 | 0.000777778 | 2.524113178 |
| Frl | Q9VUC6;Q9VUC6-2;Q9VUC6-3;Q9VUC6-4 | + | 5.563669418 | 0 | 2.516463757 |
| anon-15Db | Q9VX80 | + | 3.374847504 | 0.001327869 | 2.510277271 |
| BcDNA:AT13896 | Q0KI69;Q9VFA3 | + | 3.579682991 | 0.001241379 | 2.505836725 |
| gek | Q9W1B0 | + | 3.080514772 | 0.002728324 | 2.498746395 |
| Rbpn-5 | Q9W2M0 | + | 2.684923323 | 0.004508539 | 2.496180058 |
| Dmel\CG5886 | Q9VBL3 | + | 2.707605001 | 0.004375479 | 2.495396137 |
| RapGAP1 | A8DYW8;Q59DZ5;Q9VM00;Q9VM02 | + | 4.510567477 | 0 | 2.489897966 |
| Dmel\CG7044 | Q9VDE8 | + | 3.795270412 | 0.001113924 | 2.487280607 |
| Sarm | A8JNN2;M9NE45;M9PHR5;Q6IDD9;Q6IDD9-2 | + | 2.667994825 | 0.00471161 | 2.482356548 |
| Dmel\CG13531 | Q9W1Y8 | + | 2.635584357 | 0.005068592 | 2.475940466 |
| Dmel\CG6330 | Q9VBA0 | + | 2.317597371 | 0.009690834 | 2.474307299 |
| Zyx | Q8T0F5;Q9N675 | + | 4.341248455 | 0 | 2.472609043 |
| Clc | Q9VWA1 | + | 2.533372208 | 0.005966777 | 2.471163511 |
| dsh | P51140 | + | 4.294643802 | 0 | 2.468535185 |
| slmb | Q9VDE3 | + | 4.759263962 | 0 | 2.466529846 |
| Vps8 | Q9VRX2 | + | 2.343485491 | 0.009411437 | 2.461975574 |
| Dmel\CG7139 | Q9XZ11 | + | 3.058531537 | 0.002805634 | 2.459967613 |
| l(3)04053 | Q9VNR6 | + | 3.559982371 | 0.001229268 | 2.454724789 |
| Orai | Q9U6B8-3 | + | 2.778595541 | 0.004 | 2.450080156 |
| Dmel\CG10435 | Q9VHX1 | + | 2.747375969 | 0.004144 | 2.448090792 |
| Cka | D3DML3;M9NEC0;M9PF42;Q9VLT9 | + | 3.80429894 | 0.001025641 | 2.444633722 |
| Unc-115a;Unc-115b | A0A0B4LGY1;A0A0B4LGZ0;A0A0B4LI11;E1JIH2;E1JIH3;Q8INN6;Q9VH91;Q9VH92 | + | 3.498177406 | 0.00132093 | 2.437742949 |
| cmb | Q9VU76 | + | 2.806934376 | 0.003881104 | 2.431368589 |
| RtGEF | A8DZ20 | + | 2.643001988 | 0.005038391 | 2.426496983 |
| Atg17 | Q7KTS2 | + | 3.776684423 | 0.00108642 | 2.421165943 |
| Ubc6 | P25153 | + | 3.247962304 | 0.001935943 | 2.411901712 |
| baz | Q9VX75;X2JFU8 | + | 2.933433751 | 0.003182482 | 2.407409668 |
| Dmel\CG5608 | Q9VG59 | + | 2.404872356 | 0.008244898 | 2.405991793 |
| ssp2 | Q9VUA5 | + | 3.271374199 | 0.001671642 | 2.405912876 |
| Nost | X2JAX8 | + | 2.695582787 | 0.004525714 | 2.403544903 |
| Crtc | M9PFU2;Q9VVJ8 | + | 2.746354397 | 0.00412749 | 2.39794755 |
| Veneno | Q9VHR5 | + | 3.222073086 | 0.002096552 | 2.397619247 |
| Asap | A1Z7A6;A1Z7A6-2 | + | 3.104839937 | 0.00252381 | 2.395787477 |
| phtf | Q9V9A8 | + | 2.640765627 | 0.005010909 | 2.387850285 |
| Rassf | Q9VCQ7 | + | 2.462307258 | 0.007250774 | 2.387187719 |
| Ipp | Q9VK21 | + | 2.992213234 | 0.002926316 | 2.386663437 |
| dlt | Q8T626 | + | 3.830997728 | 0.000761905 | 2.383552313 |
| Cog4 | Q95TN4 | + | 2.942781828 | 0.003127451 | 2.382169962 |
| nocte | M9PE74;Q9W2U7 | + | 4.573436716 | 0 | 2.376320362 |
| milt | Q960V3;Q960V3-2;Q960V3-4 | + | 3.624826068 | 0.001109948 | 2.37595129 |
| fry | E2QCY4;M9ND72;M9NE70;M9PEP1;Q9VT28 | + | 2.923268939 | 0.003207637 | 2.375384092 |
| CenG1A | Q9NGC3;Q9NGC3-2 | + | 2.454563309 | 0.007423313 | 2.373537064 |
| Ack | Q9VZI2 | + | 4.078254027 | 0.000545455 | 2.36992836 |
| Axn | Q9V407 | + | 3.966533042 | 0.000617886 | 2.369443893 |
| Vsp37A | O77259;Q7YZA5 | + | 4.669313729 | 0 | 2.366384029 |
| Vps16B | Q9VAG4 | + | 2.709490681 | 0.004323077 | 2.365203142 |
| Pfdn6 | Q9VW56 | + | 2.881044071 | 0.003316629 | 2.364881992 |
| tun | Q7K2Y9 | + | 3.573008312 | 0.001235294 | 2.35643363 |
| tamo | Q9W1A4 | + | 2.482118771 | 0.006827586 | 2.355247974 |
| Lk6 | Q9VGI4;Q9VGI5 | + | 4.253595447 | 0 | 2.352066517 |
| didum | A1Z6Z8;A1Z6Z9 | + | 2.490505327 | 0.006633071 | 2.350027084 |
| FAM21 | A1ZBW7 | + | 4.356202797 | 0 | 2.339818954 |
| Not3 | Q7K126 | + | 4.975565331 | 0 | 2.339253426 |
| CG31653;CG3805;Dm_2L:23546;Dmel\CG34126 | M9PB37;M9PC10;M9PC65;M9PEM2;Q8IPL3 | + | 3.317630907 | 0.001407115 | 2.33690238 |
| Trpm | A8DYE2;A8DYE2-4 | + | 2.989791279 | 0.002895833 | 2.336657524 |
| Sik3 | A0A0B4LFW1;A1ZBC4 | + | 4.180034108 | 0.000196078 | 2.334041595 |
| Magi | Q9W2L2 | + | 2.506809282 | 0.00616129 | 2.333956242 |
| CG11846 | Q9VF17 | + | 2.524330974 | 0.006029557 | 2.333920717 |
| Not1 | A8DY80;A8DY81;E2QCN4 | + | 3.322692747 | 0.001418327 | 2.327168465 |
| aux | A0A0B4KF57;A0A0B4KG14;Q9VMY8 | + | 3.132345227 | 0.002299694 | 2.324302197 |
| CG9951 | Q9VVB4 | + | 2.437013394 | 0.007681682 | 2.321736813 |
| Tsc1 | Q9VCC9 | + | 3.200861897 | 0.002019934 | 2.318632841 |
| Tmf | Q9W3V2 | + | 2.758001842 | 0.004129555 | 2.298424482 |
| mon2 | Q9VLT1 | + | 3.237251556 | 0.002021053 | 2.291716337 |
| Girdin | M9PDY5 | + | 2.784458942 | 0.003983402 | 2.285425425 |
| Cnot4 | M9MRH8;M9PB92 | + | 2.428342035 | 0.007748879 | 2.28502655 |
| Dmel\CG11168 | A0A0B4KH07;Q9VC09 | + | 2.89140829 | 0.00324424 | 2.284942865 |
| Lasp | Q8I7C3 | + | 3.00793505 | 0.00286631 | 2.284285784 |
| l(3)05822 | Q9VE96 | + | 3.706007903 | 0.001211429 | 2.271706581 |
| Wnk | M9PGC5;M9PGC5-2 | + | 3.843202482 | 0.000671329 | 2.267282248 |
| Golgin245 | Q9W1R3 | + | 2.7237013 | 0.004116279 | 2.263256311 |
| mop | Q9VUH6 | + | 2.698895558 | 0.004497132 | 2.262451172 |
| retm | Q9VMD6 | + | 3.587746926 | 0.001247525 | 2.260636091 |
| Ge-1 | Q9VKK1 | + | 2.636433115 | 0.005077758 | 2.260054588 |
| Edc3 | Q9VVI2 | + | 2.881790145 | 0.003287671 | 2.254445553 |
| SKI3 | Q6NNB2 | + | 3.262300475 | 0.001647059 | 2.254287243 |
| Mhcl | A0A0B4KGR6;Q0KI67;Q8INC1 | + | 3.417897012 | 0.001327434 | 2.252527952 |
| Rok | Q9VXE3;X2JC83;X2JKQ8 | + | 2.814147365 | 0.003853448 | 2.251194477 |
| Gyf | Q7KQM6;Q7KQM6-2 | + | 3.776805663 | 0.001093168 | 2.250991821 |
| Pez | Q9V3V3 | + | 3.878335027 | 0.000716418 | 2.250920773 |
| hpo | Q8T0S6 | + | 4.314604414 | 0 | 2.249811172 |
| CG18490;CG34240 | M9PF53;Q9VTH2;X2J8Y9 | + | 3.277692083 | 0.0016 | 2.24739337 |
| Eps-15 | Q8MMD2;Q8MMD3 | + | 2.827729821 | 0.003817391 | 2.229768991 |
| Pkn | A1Z7T0;A1Z7T0-2;A1Z7T1;A1Z7T3;A8DY76 | + | 2.954783829 | 0.003024876 | 2.22884798 |
| Wdr92 | Q9VVM7 | + | 2.929098882 | 0.003246377 | 2.227421522 |
| Liprin-alpha | M9PCD4;M9PCR7;Q9VM93 | + | 2.586717589 | 0.005459552 | 2.215614319 |
| 4E-T | Q8IH18;Q8IH18-2 | + | 3.40219528 | 0.001390558 | 2.209429264 |
| DCP1 | Q9W1H5 | + | 2.381067402 | 0.008670487 | 2.20473671 |
| Ccz1 | Q9VZL5 | + | 4.776376471 | 0 | 2.204118013 |
| IPIP | Q9VMI6 | + | 2.64332686 | 0.005047619 | 2.19595027 |
| CG5521 | Q9VB98;Q9VB98-2 | + | 2.599713938 | 0.005302977 | 2.192576647 |
| Gprk1 | P32865 | + | 2.595887562 | 0.005275261 | 2.192163467 |
| msps | A0A0B4K664;A0A0B4K700;Q9VEZ3 | + | 3.198293473 | 0.002006601 | 2.190027714 |
| CG6729 | Q9VKQ6 | + | 5.329096018 | 0 | 2.18995738 |
| Dmel\CG3632 | A8JV04;Q7YU03 | + | 3.384584731 | 0.00135 | 2.189929008 |
| DEF8 | Q9VTT9 | + | 2.330961328 | 0.009465565 | 2.188343525 |
| Dmel\CG5056 | Q9VKY0 | + | 2.997009679 | 0.002914894 | 2.187653065 |
| Aven | Q9VYK8 | + | 2.85958642 | 0.003482143 | 2.185645819 |
| lqfR | Q8IN05;Q9VD13 | + | 3.486673973 | 0.001308756 | 2.185128212 |
| Dmel\CG8060 | A1ZAE2 | + | 2.726074203 | 0.004124272 | 2.183282614 |
| X11L | E1JJN6;Q9VX41;X2JE74 | + | 5.234156882 | 0 | 2.181540966 |
| GckIII | Q9VEN3 | + | 2.941694154 | 0.003119804 | 2.180987358 |
| CG13768;CT33252 | A8DYW2;Q9VMA9 | + | 3.140268231 | 0.002149068 | 2.179249048 |
| cyst | Q9VIV0 | + | 3.713025617 | 0.001239766 | 2.177959919 |
| kri | M9PHI7;Q9VRQ1 | + | 4.680172809 | 0 | 2.172186852 |
| Ugt37D1 | Q9VJH8 | + | 2.324672884 | 0.009454047 | 2.169992447 |
| Chc | P29742 | + | 3.196646741 | 0.002 | 2.168961048 |
| tapas | D5AEP3;Q0E910 | + | 4.27429645 | 0 | 2.164254904 |
| p115 | Q9W3N6 | + | 2.304469766 | 0.00994844 | 2.161016226 |
| CG8202 | Q9VHM2 | + | 2.893678678 | 0.003214781 | 2.159109592 |
| Atg13 | Q9VHR6 | + | 3.668106148 | 0.00115847 | 2.148596525 |
| Afti | A0A4D6K2J4;Q9VFK5 | + | 3.261317354 | 0.001641026 | 2.147226334 |
| fmt | M9PEB1;Q9VRV1 | + | 2.582371206 | 0.005553265 | 2.146846533 |
| Apc;Apc2 | A0A1Z1CN92;Q9VAS9;Q9Y1T2 | + | 4.597733042 | 0 | 2.14500165 |
| Dmel\CG8578 | Q9VXN3 | + | 2.717239724 | 0.004177606 | 2.143606186 |
| subdued | A0A0B4JCY1;Q9VDV4 | + | 3.127699362 | 0.002339394 | 2.136855125 |
| hook | Q24185 | + | 2.415629543 | 0.008041237 | 2.136849165 |
| Dmel\CG4853 | A0A0B4LFJ8;A1ZAU6 | + | 4.42855978 | 0 | 2.135308027 |
| Kaz | Q8MSB1 | + | 3.165476329 | 0.002101911 | 2.135253906 |
| numb | P16554;P16554-2 | + | 2.673351696 | 0.004669173 | 2.133548498 |
| wmd | Q9W1N4 | + | 4.452922002 | 0 | 2.127949238 |
| GM10035 | A8Y582;Q7PLP4;Q7PLP5 | + | 4.306548831 | 0 | 2.125643492 |
| RhoGEF2 | A0A0B4K7T7;A1ZAN6 | + | 3.833754859 | 0.000772414 | 2.124756813 |
| Dmel\CG10283 | E1JHK9;Q9VJ84 | + | 6.455373737 | 0 | 2.123837948 |
| cnn | P54623-3 | + | 2.973639449 | 0.002946292 | 2.122824907 |
| Atg1 | Q8MQJ7 | + | 3.807255451 | 0.001038961 | 2.122025251 |
| PAPLA1 | Q9VLS7;X2JDJ3 | + | 2.511829714 | 0.006156352 | 2.120210409 |
| POSH | Q7K4D1 | + | 4.292395824 | 0 | 2.119302511 |
| Arp3 | P32392 | + | 2.777113801 | 0.003983573 | 2.113761902 |
| Wdr59 | Q9VKK2 | + | 4.289870764 | 0 | 2.113741875 |
| Dmel\CG6005 | Q9VE12 | + | 3.644854193 | 0.001145946 | 2.113581896 |
| NAAT1 | Q9W4C5 | + | 3.643777653 | 0.001139785 | 2.112239361 |
| cact | Q03017;Q03017-2 | + | 3.248901897 | 0.001871429 | 2.106048107 |
| Cog3 | Q961G1 | + | 2.567106746 | 0.005714286 | 2.105830669 |
| dor | Q24314 | + | 2.606737748 | 0.005224779 | 2.104815006 |
| Unr | B7Z0E2 | + | 2.972594591 | 0.002938776 | 2.101971865 |
| wash | Q7JW27 | + | 5.833356549 | 0 | 2.100337505 |
| Vps39 | Q9VEA2 | + | 2.377003836 | 0.008771429 | 2.097782612 |
| loco | Q9VCX1;Q9VCX1-3 | + | 5.949892611 | 0 | 2.095741749 |
| Rilpl | O76878 | + | 2.996202343 | 0.002907162 | 2.09564209 |
| PhKgamma | Q961E7;Q9I7D0 | + | 3.3823561 | 0.001338843 | 2.095547676 |
| r2d2 | Q9VLW8 | + | 2.438215077 | 0.007704819 | 2.092665195 |
| PTP-ER | Q9W2F3 | + | 2.516682247 | 0.006039216 | 2.08680582 |
| Naus | Q8SX68 | + | 4.008117275 | 0.000644068 | 2.084895611 |
| wts | Q9VA38 | + | 2.657721933 | 0.004945996 | 2.082135916 |
| Dmel\CG6688 | Q9VCS4 | + | 2.437652607 | 0.007693233 | 2.081475258 |
| btsz | A0A0B4K657;A8JR10;B7Z0K2;Q6XK19 | + | 2.812450809 | 0.003820513 | 2.07951808 |
| yki | Q45VV3 | + | 6.140517884 | 0 | 2.073634148 |
| Vps51 | Q8MSY4 | + | 2.41580191 | 0.008053097 | 2.069259644 |
| cv-2 | Q9W2H2 | + | 2.961911345 | 0.003015075 | 2.069074154 |
| Dmel\CG15602 | Q4QQ70 | + | 3.696048234 | 0.001204545 | 2.06873107 |
| Sec8 | Q9VNH6 | + | 2.726941204 | 0.004132296 | 2.067472935 |
| CG15099 | A1ZBE8 | + | 3.845396979 | 0.000680851 | 2.062516928 |
| kermit | Q7JX82 | + | 2.302043152 | 0.009994609 | 2.060725927 |
| Dmel\CG9356 | Q9VHE3 | + | 2.906539447 | 0.003185012 | 2.056205988 |
| Abl | M9PCT8;M9PFI5;M9PFS1;M9PFX0;P00522 | + | 3.986535818 | 0.000628099 | 2.054785013 |
| Iml1 | Q9W0E3 | + | 3.810327827 | 0.000888889 | 2.053945303 |
| tud | P25823 | + | 2.729266042 | 0.004148438 | 2.047039747 |
| l(2)gl | P08111 | + | 3.058073834 | 0.002797753 | 2.046175957 |
| FBpp0111587 | B7Z138;Q8IRL5;Q9W2X2;X2JB32;X2JD61;X2JEC6 | + | 2.397320519 | 0.008492041 | 2.041499138 |
| betaggt-I | M9PEL6;Q24173 | + | 3.548586558 | 0.001211538 | 2.039492607 |
| CYLD | Q8IPC5 | + | 3.846209341 | 0.000685714 | 2.034888983 |
| gig | M9PD17;Q9VW83 | + | 3.775569072 | 0.001079755 | 2.03164196 |
| scrib | A0A0B4K6I1;A0A0B4K7W1;A0A0B4KI37;Q7KRY7-10;Q7KRY7-7 | + | 4.443632893 | 0 | 2.029049397 |
| mgr | Q9VGP6 | + | 3.539249271 | 0.001352381 | 2.028801203 |
| dia | P48608 | + | 3.647945947 | 0.001152174 | 2.017967939 |
| dl | P15330;P15330-2 | + | 2.663144232 | 0.00482243 | 2.014079332 |
| rut | M9PH52;P32870 | + | 5.788239678 | 0 | 2.012799978 |
| Mesh1 | Q9VAM9 | + | 4.383156263 | 0 | 2.012408257 |
| GramD1B | A8DYU1;R9PY34 | + | 3.906412602 | 0.000732824 | 2.004972935 |
| Dmel\CG11360 | Q9V4F3 | + | 2.617441118 | 0.005232975 | 2.004443645 |
| unk | Q86B79 | + | 4.032453454 | 0.000512821 | 1.99982357 |
| Ehbp1 | A1ZAN2;A4UZL2;A4UZL3;B7YZI1 | + | 2.354092336 | 0.009179775 | 1.993971586 |
| CG1846 | Q9VY47 | + | 4.803436128 | 0 | 1.989115 |
| pns | Q9W0E1 | + | 2.878153974 | 0.003363636 | 1.988580227 |
| dbo | Q9VUU5 | + | 2.928668126 | 0.003238554 | 1.983711481 |
| tyf | Q9W4M7 | + | 3.865772002 | 0.00070073 | 1.983078241 |
| TBC1D5 | Q9VFX6 | + | 4.458899147 | 0 | 1.981564522 |
| Patr-1 | Q9VEN9 | + | 2.365315557 | 0.008871287 | 1.981497288 |
| spg | A0A0B4KHD2;B7Z0R2;Q9VAS8 | + | 2.848757841 | 0.003665198 | 1.979546785 |
| l(3)76BDm | Q9VW22 | + | 2.7370535 | 0.004101961 | 1.977715969 |
| Cnb | Q9I7U5 | + | 3.240051184 | 0.002035336 | 1.977655172 |
| insc | Q9W2R4 | + | 2.951705492 | 0.003002469 | 1.97759223 |
| Nin | M9MRD3;M9PB42;M9PC27;Q8IPK8;Q9VMS6 | + | 4.628599889 | 0 | 1.976949215 |
| NP_649372 | Q9VNX0 | + | 2.493307844 | 0.00659271 | 1.975971222 |
| Dmel\CG2258 | Q9W3K6 | + | 4.761857271 | 0 | 1.969524384 |
| Sec15 | Q9VDE6 | + | 3.388187779 | 0.001355649 | 1.969150305 |
| unc-45 | Q9VHW4 | + | 2.804456612 | 0.003872881 | 1.969147444 |
| Dmel\CG4325 | Q9XZS4 | + | 4.868843853 | 0 | 1.963051081 |
| csw | P29349;P29349-2;P29349-4 | + | 3.10352251 | 0.00251632 | 1.960195303 |
| Mon1 | Q9VR38 | + | 3.411798909 | 0.001293103 | 1.959560156 |
| Dmel\CG7518 | A0A0B4K6G6;Q9VG05 | + | 4.82783199 | 0 | 1.953835964 |
| Smg5 | Q9V414 | + | 2.604311776 | 0.005197183 | 1.952928782 |
| Helz | Q9VUH9 | + | 3.202173843 | 0.002033445 | 1.952834129 |
| Sara | Q7K9H6 | + | 2.794226268 | 0.00392437 | 1.951042414 |
| Usp10 | M9PDK3;Q9W0L7 | + | 3.371731979 | 0.001322449 | 1.947560072 |
| yuri | Q8IP47;Q9V435;Q9VJP8 | + | 2.590300214 | 0.005393414 | 1.945205927 |
| Tango10 | Q7K187 | + | 2.411215497 | 0.008122987 | 1.944494486 |
| RhoGAP92B | Q9VDS5 | + | 2.907763542 | 0.003192488 | 1.943749905 |
| Dmel\CG12024 | M9MRY7;Q8IRG7;Q8SZU6 | + | 2.465949075 | 0.00725972 | 1.939339399 |
| CG7701;Dmel\CG30015 | A0A0B4KFA3;A8DY97;Q0E9C6 | + | 4.878138538 | 0 | 1.93414402 |
| Zdhhc8 | Q9W345 | + | 2.658373714 | 0.00491791 | 1.932604313 |
| Diap1 | Q24306 | + | 4.034287276 | 0.000517241 | 1.929753542 |
| obe | Q9VF56 | + | 3.127238327 | 0.002332326 | 1.927899122 |
| Ufd4 | Q9VL06 | + | 3.333246347 | 0.00136 | 1.926891565 |
| Pi3K92E | P91634 | + | 2.930725727 | 0.003254237 | 1.925328493 |
| cpa | Q9W2N0 | + | 2.827722584 | 0.003809111 | 1.914732218 |
| Pdk1 | Q9W0V1;Q9W0V1-1;Q9W0V1-2 | + | 2.303861702 | 0.00995664 | 1.912662745 |
| C3G | O77086;O77086-2;O77086-3 | + | 4.266085501 | 0 | 1.911797523 |
| Bruce | A0A0B4KH36 | + | 3.083080942 | 0.002736232 | 1.909109116 |
| ssh | Q6NN85;Q6NN85-2 | + | 3.24233352 | 0.002042553 | 1.908508062 |
| roq | Q9VV48 | + | 3.100968894 | 0.0026 | 1.908254385 |
| Clic | Q9VY78 | + | 3.629289868 | 0.001121693 | 1.906472445 |
| raptor | Q9W437 | + | 3.800197742 | 0.001121019 | 1.906063318 |
| l(1)G0196 | Q9VR59-6 | + | 2.953505365 | 0.003009901 | 1.905887842 |
| Pfdn2 | Q9VTE5 | + | 2.495416433 | 0.006505564 | 1.904630661 |
| cg12129 | Q7JW66 | + | 5.212863145 | 0 | 1.902220726 |
| tst | Q9VCH8 | + | 2.432791347 | 0.007760479 | 1.899905682 |
| Dph2 | Q9VFE9 | + | 4.05298614 | 0.000540541 | 1.89697814 |
| Use1 | Q9VSU7;Q9VSU7-2 | + | 2.780828482 | 0.004016563 | 1.896116972 |
| Gga | Q9W329 | + | 2.493204946 | 0.006582278 | 1.892036915 |
| boca | Q8T9B6 | + | 3.006862555 | 0.002858667 | 1.889627218 |
| Arpc2 | Q9VIM5 | + | 3.037335093 | 0.002759003 | 1.88888073 |
| par-1 | A1ZBL7;E1JGN0;Q6NPA6 | + | 3.708296513 | 0.001225434 | 1.887466431 |
| fwd | A0A4D6K881;Q9W0R8 | + | 4.080101073 | 0.000550459 | 1.883631706 |
| Snx17 | Q9VL28 | + | 2.918892047 | 0.003184834 | 1.883368254 |
| Usp8 | Q9VDD8 | + | 2.812636245 | 0.003828694 | 1.87962389 |
| elgi | Q9VUV7 | + | 3.061599809 | 0.00282153 | 1.877015591 |
| Cog1 | Q9VGC3 | + | 2.332684202 | 0.009482711 | 1.875427008 |
| Klp98A | Q9VB25 | + | 2.300834892 | 0.009994624 | 1.874911308 |
| SMAP | A1Z7K6 | + | 2.644291513 | 0.005056881 | 1.872730017 |
| vlc | A1Z6H4;Q7K1T1 | + | 2.859569882 | 0.003474388 | 1.867072344 |
| hep | Q23977-3 | + | 3.009286627 | 0.002873995 | 1.866709948 |
| p120ctn | Q7PLI0 | + | 3.062148394 | 0.002837607 | 1.866455078 |
| Sou | Q9W2N5 | + | 2.449501089 | 0.007409437 | 1.865507841 |
| ck | Q9V3Z6 | + | 3.064698935 | 0.002845714 | 1.863633394 |
| Vps13B | Q9VAD3 | + | 2.898893469 | 0.003244755 | 1.8611238 |
| pigs | Q9W3Y4 | + | 2.461538723 | 0.007228395 | 1.860713243 |
| Arp2 | P45888-2 | + | 2.393864454 | 0.008560694 | 1.859340429 |
| Dmel\CG11710 | Q8T921;Q9VRA1 | + | 5.207855 | 0 | 1.858958721 |
| jub | Q9VY77 | + | 3.851683863 | 0.000695652 | 1.858669758 |
| CG13811 | B7Z087;B7Z088;M9PFQ9 | + | 3.670911041 | 0.001171271 | 1.857579947 |
| Wdr24 | Q9XZ25 | + | 2.526202488 | 0.006009885 | 1.85640645 |
| Khc-73 | A0A0B4KFR5;A0A0B4LGK5;A4UZI0 | + | 3.216794543 | 0.002068027 | 1.854574203 |
| Atx2 | Q8SWR8;Q8SWR8-2;Q8SWR8-3 | + | 3.317065865 | 0.001396078 | 1.853031635 |
| Vrp1 | A0A0B4LH48;A8DYL3;E1JGR7;E1JGR9;Q0E8Y8;Q7KVL6;Q8SX98 | + | 2.495781916 | 0.006477707 | 1.851508379 |
| Epg5 | Q9VE34 | + | 2.873218315 | 0.003340858 | 1.848829269 |
| Liprin-beta | Q9VU88 | + | 2.345985311 | 0.009383754 | 1.848515272 |
| Arp10 | Q9VWE8 | + | 2.441235677 | 0.007637462 | 1.846232176 |
| CG5340 | Q86B82;Q86B83 | + | 3.635207159 | 0.00113369 | 1.844907999 |
| Tbce | A1Z6J5 | + | 2.600119462 | 0.005270175 | 1.842107534 |
| raskol | M9MSE8;Q8IQZ6;Q8T498;Q8T498-2;X2JCD6;X2JFL7 | + | 2.873223392 | 0.003348416 | 1.839778662 |
| chb | Q9NBD7 | + | 2.714793693 | 0.004200385 | 1.838378906 |
| DmTOM1 | M9PF36;Q9VSZ1 | + | 2.588202879 | 0.005406897 | 1.830591917 |
| Dmel\CG1344 | A1Z6G6 | + | 3.042319797 | 0.002774373 | 1.824459314 |
| pnut | P40797 | + | 2.887816838 | 0.003258581 | 1.822873116 |
| Src64B | P00528 | + | 2.656422871 | 0.004927644 | 1.82143259 |
| Sec10 | Q9XTM1 | + | 2.302756579 | 0.009975709 | 1.820799351 |
| Mical | Q86BA1 | + | 4.67311633 | 0 | 1.817554951 |
| Dmel\CG8671;anon-WO0118547.182 | A2VEG3;M9PDK6;M9PDX3;Q9VID5 | + | 2.727837087 | 0.004140351 | 1.812645197 |
| stv | A8JNS4;M9PCD9;Q9VU81;Q9VU82;Q9VU83 | + | 2.576483101 | 0.005684932 | 1.810697079 |
| Pngl | Q7KRR5 | + | 3.025419724 | 0.002757493 | 1.809933662 |
| Hsc70-1;Hsc70-2;Hsc70-3;Hsc70-4;Hsp68 | O97125;P11146;P11147;P29843;P29844 | + | 2.919985376 | 0.0032 | 1.805874825 |
| Dmel\CG16952 | A0A4D6K4K0;Q9VXM6;X2JFM8 | + | 3.748234056 | 0.001053892 | 1.805726528 |
| ric8a | Q9W358 | + | 2.301937845 | 0.009981157 | 1.804832935 |
| Pi3K21B | Q7KTZ2 | + | 4.747166104 | 0 | 1.802154779 |
| mTor | Q9VK45 | + | 2.988848281 | 0.002888312 | 1.80150485 |
| Tnpo | Q9VRV8 | + | 3.355971703 | 0.001365462 | 1.800500393 |
| Dcr-2 | A1ZAW0 | + | 2.529554516 | 0.00594702 | 1.800333261 |
| Ankle2 | Q8MQX9;Q8MQX9-2;Q8MQX9-3 | + | 3.139943665 | 0.002142415 | 1.798039198 |
| Tbc1d15-17 | Q9VPL5 | + | 2.99104849 | 0.002910995 | 1.794663668 |
| Lst8 | Q9W328 | + | 2.471203852 | 0.00715 | 1.790049791 |
| Rme-8 | A0A0B4LFW6;A1Z7S0 | + | 3.177005336 | 0.00207074 | 1.788804054 |
| Pak3 | A0A0B4KGS4;Q9VEV1 | + | 3.250226586 | 0.001827338 | 1.785252333 |
| tral | M9PFF9;M9PFG3 | + | 2.534174374 | 0.005976705 | 1.784784794 |
| Atg16 | B7Z0R7;Q86BR6 | + | 2.615832503 | 0.005242857 | 1.783429384 |
| ArfGAP3 | A8JNX0;M9PG88;M9PGG0;M9PIL3;Q9VNS2 | + | 2.946286956 | 0.003027027 | 1.775939941 |
| CG11945 | Q9VCH2 | + | 2.946343528 | 0.003034483 | 1.775020838 |
| Dmel\CG1407 | A0A0B4LEL6;A0A0B4LF00;A0A0B4LF16;A1Z833 | + | 3.936421159 | 0.00059375 | 1.77306366 |
| poe | Q9VLT5 | + | 2.647916852 | 0.005023941 | 1.771598101 |
| Ced-12 | Q9VKB2 | + | 3.596898326 | 0.001174129 | 1.770684958 |
| cindr | A0A0B4KI34;Q9VA36 | + | 2.991972928 | 0.002918635 | 1.767367363 |
| krz | Q9V393 | + | 2.617826893 | 0.00524237 | 1.763982534 |
| garz | A1Z8W8 | + | 2.606908011 | 0.005234043 | 1.7629807 |
| bbg | Q9VUE8 | + | 2.313268434 | 0.009694823 | 1.7595644 |
| Dmel\CG15744 | Q7KV24 | + | 3.400907701 | 0.001384615 | 1.759321451 |
| Snx16 | Q7JR96 | + | 4.04754291 | 0.000530973 | 1.753646851 |
| Septin1 | P42207 | + | 2.737523094 | 0.00411811 | 1.747605562 |
| Vps13D | Q9VU08 | + | 2.809413961 | 0.003889362 | 1.744052649 |
| CG8475 | Q9VLS1 | + | 3.219966187 | 0.002082192 | 1.743880987 |
| 14-3-3epsilon;14-3-3zeta | P29310;P29310-2;P29310-3;P92177;P92177-1;P92177-2;P92177-4 | + | 2.986125741 | 0.002880829 | 1.739215612 |
| Dmel\CG2747 | A0A0B4KGS8;Q32KD3;Q494I1 | + | 3.277394198 | 0.001593985 | 1.737316608 |
| LRR | A1Z734;E1JGZ8 | + | 2.888131559 | 0.003266055 | 1.736992836 |
| Vps36 | Q9VU87 | + | 3.943892954 | 0.000598425 | 1.735896349 |
| Pi3K59F | Q9W1M7 | + | 2.465486303 | 0.007248447 | 1.734733582 |
| AP-1gamma | Q7KVR7;Q7KVR8;Q86B59;Q9W388 | + | 3.027399432 | 0.002772603 | 1.733296394 |
| RhoGAPp190 | Q9VX32-2 | + | 3.906463653 | 0.000738462 | 1.727854013 |
| Dmel\CG42353 | Q9VXJ7 | + | 2.816130171 | 0.003861771 | 1.727634907 |
| sick | Q9VIQ9;Q9VIQ9-3;Q9VIQ9-4;Q9VIR0 | + | 3.383421578 | 0.001344398 | 1.726329088 |
| epsilonCOP | Q9Y0Y5 | + | 2.400031271 | 0.008261248 | 1.726133347 |
| Arpc3A | Q9VF28 | + | 3.150567886 | 0.002132492 | 1.723907471 |
| Dmel\CG12237 | Q9VWF0 | + | 3.053888389 | 0.002782123 | 1.719581842 |
| Hyccin | Q7K1C5 | + | 3.093303279 | 0.002577259 | 1.708598614 |
| msn | Q9W002 | + | 3.216940287 | 0.002075085 | 1.707610369 |
| Vps29 | Q9VPX5 | + | 2.553190039 | 0.005807757 | 1.707250118 |
| cv-c | A8JR05 | + | 2.52881314 | 0.00597686 | 1.702212334 |
| stau | P25159;P25159-2 | + | 3.825955747 | 0.000751678 | 1.701544523 |
| Diap2 | Q24307 | + | 2.614212588 | 0.005233512 | 1.700155258 |
| Dysb | Q9VVT5 | + | 3.09764618 | 0.002584795 | 1.691937447 |
| Tomosyn | Q9VYK6 | + | 2.507097737 | 0.006171244 | 1.689076185 |
| eIF4G1 | A8DZ29 | + | 2.926160129 | 0.003215311 | 1.685479164 |
| Su(fu) | Q9VG38 | + | 2.502297613 | 0.006157303 | 1.683870077 |
| Rop | Q07327 | + | 2.859753528 | 0.003489933 | 1.676234961 |
| Sbf | Q7KSP6;Q9VGH9 | + | 3.029285083 | 0.002787879 | 1.67458272 |
| PpV | Q27884 | + | 3.24996896 | 0.001878136 | 1.672930241 |
| Rga | Q94547;Q94547-2 | + | 3.985584509 | 0.000622951 | 1.672724724 |
| Dmel\CG5776 | Q9VK63 | + | 2.827133995 | 0.003800866 | 1.66873312 |
| mbc | A0A0B4K7Q6;Q9VCH4 | + | 2.478928395 | 0.006985915 | 1.666499138 |
| Dredd | Q8IRY7;Q8IRY7-2;Q8IRY7-4 | + | 3.316105017 | 0.001390625 | 1.666095257 |
| Mrtf | B7Z043;Q9VZY2 | + | 3.019555326 | 0.0028 | 1.663043261 |
| CG15612;Dmel\CG30456 | A0A0B4KEX9;A1ZAQ3 | + | 3.444924258 | 0.001279279 | 1.659580946 |
| uri | Q9W148 | + | 4.732694816 | 0 | 1.658887625 |
| lqf | A8JNM3;M9PBU6;M9PEF2;M9PEM7;Q7KU90;Q7KU93;Q9VS85 | + | 2.961410431 | 0.003007519 | 1.656019688 |
| mdcds_8484 | M9PCP3;Q9VMK9 | + | 3.122019322 | 0.002325301 | 1.650006294 |
| stck | Q8INQ9;Q8INR0 | + | 3.609456104 | 0.001087179 | 1.647035599 |
| rictor | Q9VWJ6;X2JFR5;X2JL73 | + | 3.362442005 | 0.001311741 | 1.646609783 |
| beta'COP | O62621 | + | 2.385919853 | 0.008586207 | 1.644915581 |
| stc | P40798;P40798-2 | + | 2.927335799 | 0.003223022 | 1.644536257 |
| CASK | Q24210 | + | 2.522370073 | 0.00600982 | 1.643354177 |
| Dmel\CG8915 | Q9VX63 | + | 2.85680323 | 0.003529933 | 1.634802818 |
| Irk1 | A0A0B4JCZ2 | + | 3.68459732 | 0.001191011 | 1.630382538 |
| Grp170 | O46067 | + | 2.50049259 | 0.006205128 | 1.630252361 |
| ksr | Q24171 | + | 4.267671014 | 0 | 1.622922182 |
| Strip | Q8IRD5;Q9VZQ9 | + | 2.778758365 | 0.004008264 | 1.611613989 |
| Dmel\CG6607 | Q9VC66 | + | 5.036670477 | 0 | 1.610354662 |
| Crk | Q9XYM0 | + | 5.978284847 | 0 | 1.607317686 |
| CG13249;CG32428-RA | C0PV00;Q9VPE6 | + | 2.490673109 | 0.006599369 | 1.604761839 |
| ssp3 | Q9VJ35;X2J6U8 | + | 2.674613533 | 0.004677966 | 1.599298716 |
| Syx16 | Q9VR90 | + | 3.869905708 | 0.000705882 | 1.598374367 |
| Fic | Q8SWV6 | + | 2.355992786 | 0.009183099 | 1.597427607 |
| CG15869;CG5100 | Q9VPD7;Q9VPE1 | + | 3.282336067 | 0.001612167 | 1.592171431 |
| mio | Q9VQ89 | + | 5.330643975 | 0 | 1.589247704 |
| Usp30 | Q9W462 | + | 2.5963378 | 0.005284468 | 1.588990927 |
| Kat60 | A0A0B4KFL3;Q9VN89 | + | 2.60880827 | 0.005243339 | 1.586363792 |
| Dmel\CG7766 | M9NDS5;M9NGW5 | + | 3.090118021 | 0.002616279 | 1.585219383 |
| spri | Q8MQW8-6 | + | 2.919536898 | 0.003192399 | 1.584827662 |
| CG9062 | Q1LZ08 | + | 2.748718884 | 0.004152305 | 1.581469059 |
| Snap29 | Q9W1I8 | + | 3.833539948 | 0.000767123 | 1.580305815 |
| Tbc1d8-9 | M9PIF1;Q9VP46 | + | 2.503669482 | 0.006167203 | 1.575004578 |
| RhoGAP15B | Q0KHR5 | + | 3.599049757 | 0.001065327 | 1.573933363 |
| dock | Q8IPW2;Q9VPU1 | + | 3.783795015 | 0.0011 | 1.570589066 |
| NAT1 | A1Z968 | + | 2.785019873 | 0.00395842 | 1.564962387 |
| Cen | Q9VIK6 | + | 4.324324133 | 0 | 1.560453176 |
| CG18638 | Q9VTV7 | + | 2.743571741 | 0.004111111 | 1.560120821 |
| CaBP1 | Q9V438 | + | 3.214653838 | 0.002061017 | 1.559159279 |
| Smurf | Q9V853 | + | 3.027635312 | 0.00278022 | 1.558123589 |
| Crag | Q9W3D3;Q9W3D3-2 | + | 3.733466053 | 0.001254438 | 1.550928593 |
| eIF4E1 | P48598;P48598-2 | + | 2.379517843 | 0.008732475 | 1.549337626 |
| mv | Q9W060 | + | 2.97161333 | 0.002931298 | 1.546806335 |
| dx | Q23985 | + | 3.81757348 | 0.000741722 | 1.545800924 |
| Dmel\CG1951 | Q9VAR0 | + | 3.830110869 | 0.000756757 | 1.543323994 |
| kappaB-Ras | Q9V4L4;Q9V4L4-2 | + | 2.435245271 | 0.007748126 | 1.540806293 |
| Tango11 | Q961C9 | + | 2.522726595 | 0.006019672 | 1.540747643 |
| Synj | Q5U0V7 | + | 2.797081944 | 0.003932632 | 1.54071331 |
| kek5 | Q9VWI6 | + | 2.452490375 | 0.007400612 | 1.537558556 |
| Dmel\CG4839 | Q9VL34 | + | 2.956831375 | 0.00304 | 1.534480572 |
| Pfdn4 | Q9VRL3 | + | 3.610347461 | 0.001092784 | 1.533056736 |
| shv | Q9VPQ2 | + | 2.52531663 | 0.006 | 1.532788038 |
| Vav | Q9NHV9 | + | 3.81293776 | 0.000894737 | 1.524074316 |
| Btk | P08630 | + | 3.364120676 | 0.001317073 | 1.522420406 |
| bun | Q24523-1 | + | 2.963511843 | 0.00302267 | 1.521568775 |
| Dmel\CG6891 | Q8IQX5;Q9VWQ7 | + | 3.016173209 | 0.002792453 | 1.520319462 |
| Wdr81 | Q9VKD0 | + | 3.466756897 | 0.001290909 | 1.519670248 |
| Kcmf1 | Q95RX5 | + | 3.252483956 | 0.001782609 | 1.519346714 |
| PDZ-GEF | B7Z025;B7Z026;Q9VMF3 | + | 3.173204766 | 0.002064103 | 1.515021801 |
| Dmel\CG13124 | Q9VL73 | + | 3.159512331 | 0.002095238 | 1.513131142 |
| Dcr-1 | Q9VCU9 | + | 2.87089968 | 0.003369369 | 1.510981321 |
| TSG101 | Q9VVA7 | + | 3.471072132 | 0.001296804 | 1.50828433 |
| Dmel\CG10011 | Q9VAU5 | + | 2.953784479 | 0.00301737 | 1.506238937 |
| Vps37B | Q9VN88 | + | 3.118525839 | 0.002426426 | 1.505154371 |
| Sema5c | Q9VTT0;Q9VTT0-2 | + | 2.412522685 | 0.008111437 | 1.5040977 |
| mtm | Q9VMI9 | + | 2.64545366 | 0.005014706 | 1.500837564 |
| Dmel\CG2701 | O76912 | + | 3.303164062 | 0.00151938 | 1.496126413 |
| slpr | Q95UN8 | + | 4.488804312 | 0 | 1.483389378 |
| Slip1 | Q8MR31 | + | 2.737488208 | 0.00411002 | 1.483060122 |
| Nha1 | Q7KTL6;Q9VM74 | + | 3.70686151 | 0.001218391 | 1.482806683 |
| Mkrn1 | Q9VP20 | + | 2.497784129 | 0.006341308 | 1.478286982 |
| PIP4K | Q8SXX1 | + | 3.412812421 | 0.001304348 | 1.477161169 |
| Dmel\CG12016 | Q9VZR0 | + | 2.982893059 | 0.00296144 | 1.476105452 |
| rush | O76902 | + | 2.757543609 | 0.004145161 | 1.475002766 |
| Txl | Q9VRP3 | + | 2.891103677 | 0.003236782 | 1.474813938 |
| HDAC4 | M9NEF2;M9PHQ2;Q9VYF3 | + | 4.328049269 | 0 | 1.473849535 |
| hop | Q24592 | + | 2.576182675 | 0.005675214 | 1.473049641 |
| Atox1 | M9PD88;Q95RR1 | + | 3.030965713 | 0.00279558 | 1.471639395 |
| Acbp2 | P42281 | + | 4.476137892 | 0 | 1.469654322 |
| CdGAPr | E1JHM0;E1JHM1;Q9VIS1 | + | 3.203725027 | 0.002040268 | 1.468760967 |
| sktl | Q9W2R3 | + | 2.344537486 | 0.009393007 | 1.467734814 |
| vap | Q8IR23 | + | 3.914020316 | 0.000744186 | 1.467627048 |
| CG5316 | Q8MSG8;Q8MSG8-2 | + | 2.368841871 | 0.008883853 | 1.465715647 |
| anon-WO0172774.52 | Q9VTY5 | + | 4.135386498 | 0.000192308 | 1.45998311 |
| Nak | Q9VJ30 | + | 3.548577645 | 0.001205742 | 1.455079079 |
| Gyc76C | Q7JQ32 | + | 2.375009615 | 0.008756757 | 1.45229578 |
| Dmel\CG2321 | Q9VAL2 | + | 2.641566316 | 0.005020036 | 1.448713303 |
| Efa6 | E1JIT7;E1JIT7-2;E1JIT7-3;E1JIT7-4;E1JIT7-5 | + | 3.683846567 | 0.001184358 | 1.448435783 |
| pins | Q9VB22 | + | 3.670467251 | 0.001164835 | 1.446903706 |
| twin | Q7K112;Q8IMX1;Q9VCB6 | + | 2.848674535 | 0.003657143 | 1.444622755 |
| FCHo2 | Q9VHC4 | + | 2.385971298 | 0.008598561 | 1.443430901 |
| Pi3K68D | Q7K3H0;Q95RR5 | + | 2.491294339 | 0.006609795 | 1.441553116 |
| pst | Q8IQ80;Q8IQ82 | + | 3.722322077 | 0.001247059 | 1.434632778 |
| Arpc4 | Q9VMH2 | + | 2.442779793 | 0.007563636 | 1.432101488 |
| CG2182 | Q9VNG1 | + | 2.314346632 | 0.009699454 | 1.429077148 |
| Mbs | A0A6M3Q7U2;A0A6M3Q913;A8JNT6;M9NDS8;Q9VUX7 | + | 2.306761971 | 0.009812245 | 1.428730249 |
| CG31035;CG7814;Dmel\CG34133 | A0A0C4DHB8;Q0KHZ4;Q8IMK1 | + | 2.453450358 | 0.007411945 | 1.42722559 |
| Smg6 | Q9VBZ6 | + | 4.244010219 | 0.00020202 | 1.412238121 |
| yrt | A0T1Z4;A0T1Z5;Q9VFU8 | + | 2.859167068 | 0.003466667 | 1.411872387 |
| lap | A0A0B4KF90;A0A0B4KFE2;A0A0B4KGC6;B7Z0U7;E1JJ78;Q9VI75 | + | 2.595772691 | 0.005266087 | 1.410616398 |
| dRNF34 | A1Z971 | + | 2.361679167 | 0.009079096 | 1.409711123 |
| sra | Q9XZL8 | + | 2.340239375 | 0.009427778 | 1.408330679 |
| Elp3 | Q9VQZ6 | + | 2.593705397 | 0.005291667 | 1.407744884 |
| FIG4 | Q9VR06 | + | 2.509494065 | 0.006175325 | 1.404124498 |
| MICAL-like | Q9VU34 | + | 2.650873491 | 0.00496679 | 1.402295113 |
| MAPk-Ak2 | P49071 | + | 4.110036508 | 0.000560748 | 1.396897316 |
| Tap42 | Q9VJZ5 | + | 2.65271033 | 0.00497597 | 1.39481473 |
| pll | Q05652 | + | 3.020639969 | 0.002807588 | 1.394597769 |
| CG6448 | M9PDE6;R9PY70 | + | 2.558193871 | 0.005827411 | 1.387313604 |
| CG12562 | Q9VP02 | + | 3.130425577 | 0.002285714 | 1.3853724 |
| anne | A8DZ26;L0MLL1;Q7KQN3;Q8IMA6 | + | 2.761812774 | 0.004137931 | 1.378898382 |
| Pten | Q7KMQ6;Q9V3L4 | + | 4.597890085 | 0 | 1.367918491 |
| Dmel\CG14647 | B9A0N7 | + | 5.04235956 | 0 | 1.367696285 |
| wake | A0A0B4K6D0;A0A0B4K6W6;A0A0B4LHH4;Q9VCW7;X4YX01 | + | 2.402847982 | 0.008285298 | 1.362947941 |
| PH4alphaSG1 | Q9VA63 | + | 3.76230228 | 0.001066667 | 1.361837387 |
| CG13284-RB;Dmel\CG13284 | D5SHN1;Q86BQ3;Q8IGQ3 | + | 2.985012227 | 0.002865979 | 1.36130023 |
| RnrS | P48592 | + | 2.459179016 | 0.007268817 | 1.355436087 |
| Dsor1 | Q24324 | + | 2.642434083 | 0.005029197 | 1.354038715 |
| Mekk1 | Q8MSQ4 | + | 4.749240742 | 0 | 1.353794336 |
| scat | Q9VLC0 | + | 2.812794403 | 0.00383691 | 1.353736877 |
| Dmel\CG5126 | Q9VPY7 | + | 2.623909245 | 0.005136691 | 1.348500252 |
| Dmel\CG6454 | A0A0B4KGY2;Q8IMW9;Q9VC89 | + | 2.901689356 | 0.00317757 | 1.347279787 |
| BEST:GH09876 | Q9VJ22 | + | 2.81386632 | 0.003845161 | 1.344332695 |
| eIF4B | Q7PLL3 | + | 3.102086111 | 0.002508876 | 1.342040539 |
| Klp64D | Q9VRK9 | + | 2.509615853 | 0.006146341 | 1.339525223 |
| strat | Q9VLP3 | + | 2.441452851 | 0.007649017 | 1.333459139 |
| Klp10A | Q960Z0 | + | 3.299841309 | 0.00157529 | 1.326741219 |
| Dmel\CG5726 | Q7JRH5 | + | 2.657148373 | 0.004936803 | 1.315307617 |
| Hn | P17276 | + | 2.303159474 | 0.009943166 | 1.314494133 |
| Vps35 | Q7KVL7;Q9W277 | + | 2.538867035 | 0.005936561 | 1.313464642 |
| Slik | B7YZQ1;Q9W179 | + | 2.757660367 | 0.004153535 | 1.311011791 |
| Nedd4;Su(dx) | Q9VVI3-2;Q9VVI3-3;Q9Y0H4 | + | 3.296990192 | 0.001630769 | 1.31041646 |
| mbt | Q9VXE5 | + | 2.417300924 | 0.008029542 | 1.305703402 |
| rl | P40417 | + | 3.023782307 | 0.002815217 | 1.304049969 |
| CG3558 | Q9VQK0;Q9VQK0-2;Q9VQK0-3 | + | 3.133385717 | 0.002306748 | 1.299722195 |
| cdm | Q9VEC5 | + | 3.186656557 | 0.001974026 | 1.298352242 |
| Eip63E | M9PE01;Q7KM03;Q7KM04;Q7KM05;Q7KM08;Q9XTK9 | + | 3.204853894 | 0.002047138 | 1.297756672 |
| Dmel\CG3744 | Q9VC19;Q9VC20 | + | 3.138412451 | 0.002135802 | 1.297727585 |
| CG7694 | Q9VE61 | + | 2.849765137 | 0.003681416 | 1.297564268 |
| Dmel\CG9205 | Q9W0K9 | + | 3.557932506 | 0.001217391 | 1.296537399 |
| Uvrag | Q9VK07 | + | 3.886536785 | 0.000721805 | 1.295546532 |
| anon-69Ag | Q9VTX1 | + | 5.390245594 | 0 | 1.292430401 |
| Fibp | M9PFY4;Q8WR19;Q9VW45 | + | 2.419682547 | 0.007988148 | 1.288633347 |
| hppy | A0A0B4LFQ3;A0A0B4LFV0;A0A0B4LGY8;A1ZBH7;A1ZBH8 | + | 3.486146266 | 0.001302752 | 1.287525177 |
| Dmel\CG17493 | M9PGG8 | + | 3.606926134 | 0.001076142 | 1.283096075 |
| Atg6 | Q9VCE1 | + | 4.125383304 | 0.000190476 | 1.276436329 |
| Shrm | A1Z9P3 | + | 3.615252896 | 0.001098446 | 1.273567915 |
| chico | Q9XTN2 | + | 5.07765233 | 0 | 1.265834332 |
| Dis3l2 | Q8IRJ7;Q9W0T7 | + | 2.500235318 | 0.0062656 | 1.265588999 |
| htt | Q9V3N4 | + | 2.450518123 | 0.007378049 | 1.262910128 |
| anon-18DEc | Q9VWD5 | + | 4.038441619 | 0.000526316 | 1.261272192 |
| MKP-4 | Q9VWF4 | + | 3.537148632 | 0.001345972 | 1.25502491 |
| ArfGAP1 | M9PFF1;Q9VTX5 | + | 3.417735276 | 0.001321586 | 1.254672527 |
| Dmel\CG8726 | A1Z782 | + | 3.430572584 | 0.001339286 | 1.253283739 |
| SH3PX1 | Q9NCC3 | + | 2.793728284 | 0.00390795 | 1.244475842 |
| Rad23 | Q9V3W9 | + | 3.417108418 | 0.001315789 | 1.243481636 |
| conu | Q8T0G4 | + | 3.168869417 | 0.002057508 | 1.224995375 |
| Pex5 | O46085;Q7KW08 | + | 2.426915158 | 0.007749627 | 1.222712994 |
| Dmel\CG6665 | A1ZAM0 | + | 2.399023052 | 0.008324638 | 1.220589876 |
| lig | A0A0B4KFC5;Q86S05-2;Q86S05-3 | + | 2.58878233 | 0.005416235 | 1.215861082 |
| Nup358 | A0A0B4K7J2;A0A0B4K7J2-2 | + | 2.639469897 | 0.005050725 | 1.215347767 |
| heca | Q9N2M8 | + | 3.412734272 | 0.001298701 | 1.212252617 |
| shep | Q8MSV2;Q8MSV2-1 | + | 2.68215908 | 0.004596226 | 1.211743355 |
| chic | P25843 | + | 3.148734424 | 0.002125786 | 1.210713387 |
| spag | Q9V3E9 | + | 3.260038657 | 0.001629091 | 1.210087061 |
| Mtmr6 | Q8MLR7;Q9W1Q6 | + | 3.184841368 | 0.001967638 | 1.207917929 |
| Csk | Q7KSQ2;Q8INJ8;Q9VGK8 | + | 2.841091309 | 0.003668122 | 1.198913574 |
| CG1161 | Q9VNA4 | + | 2.407210078 | 0.008204678 | 1.193374395 |
| Ankrd49 | Q9V3Y0 | + | 2.772341248 | 0.00396728 | 1.187497377 |
| Mad | P42003 | + | 3.147608316 | 0.002119122 | 1.186215401 |
| CG6051 | Q9VB70 | + | 2.746901139 | 0.004135729 | 1.18091774 |
| Edem2 | Q9VK27 | + | 2.767297472 | 0.004057026 | 1.177936554 |
| Med | O62609 | + | 3.709909361 | 0.001232558 | 1.170022249 |
| anon-EST:Posey170 | E1JIB4 | + | 3.599781855 | 0.001070707 | 1.166493893 |
| Cg17746 | Q9VZS1 | + | 4.768629507 | 0 | 1.163587809 |
| Atg18a | M9PEY7 | + | 2.977007274 | 0.002953846 | 1.159849644 |
| TBC1D23 | Q9VPW9 | + | 2.69862279 | 0.00448855 | 1.159267902 |
| Chmp1 | Q95SH2 | + | 2.896313147 | 0.003222222 | 1.156773329 |
| BEST:LD29996 | Q9VNA3 | + | 2.69542031 | 0.00451711 | 1.154263973 |
| sxc | Q7KJA9 | + | 2.550437726 | 0.005788235 | 1.151237011 |
| Stacl | A0A0B4KER3;A0A0B4KEV7;A0A0B4KF73;A0A0B4KFR2;A0A0B4KG20;A0A0B4LF79;A0A0B4LFG4;A0A0B4LFI1;A0A0B4LGK0 | + | 2.33062522 | 0.009452545 | 1.151016951 |
| CG3430 | Q9VM60 | + | 3.102084774 | 0.002501475 | 1.150803566 |
| Prosbeta4 | Q9VJJ0 | + | 3.056690081 | 0.002789916 | 1.144028425 |
| Atg2 | Q9VZX7 | + | 2.321044997 | 0.009523288 | 1.137847424 |
| Snrk | Q0E981 | + | 2.742408127 | 0.004094862 | 1.134452581 |
| Dmel\CG1513 | A1Z814 | + | 2.745332188 | 0.004119284 | 1.13352561 |
| ci | P19538 | + | 3.111670161 | 0.002419162 | 1.121350527 |
| CG2614 | Q9VIK9 | + | 3.222993583 | 0.002103806 | 1.118091583 |
| Stat92E | Q24151;Q24151-2 | + | 3.187456522 | 0.001993443 | 1.108749866 |
| Dmel\CG2812 | Q9W1G4 | + | 3.688455936 | 0.00119774 | 1.104408264 |
| CG-6428 | Q9W4N6 | + | 2.549163194 | 0.005825503 | 1.103816748 |
| step | Q0E8N2 | + | 3.132003345 | 0.002292683 | 1.09553194 |
| Mondo | Q8INT6;Q9VID4 | + | 2.539874696 | 0.005946488 | 1.093595028 |
| CHMP2B | Q9VRJ5 | + | 2.37540702 | 0.008769231 | 1.091192722 |
| Hou | Q9VWS1 | + | 2.76440676 | 0.004081301 | 1.089471102 |
| Dmel\CG1582 | Q9VZ55 | + | 2.332478792 | 0.009469613 | 1.088855028 |
| Mkk4 | O61444 | + | 3.026586641 | 0.002765027 | 1.075715542 |
| Dmel\CG15445 | Q9VRF3 | + | 3.133798764 | 0.002313846 | 1.071098566 |
| AP-1sigma | A0A0B4KHB7;B8A403;Q9VCF4 | + | 2.793789249 | 0.003916143 | 1.068732262 |
| su(r) | Q9W374 | + | 2.495304711 | 0.006495238 | 1.067741394 |
| CG17486 | Q5LJP9 | + | 4.539924393 | 0 | 1.061326981 |
| heph | A0A0B4K6P4;A0A0B4K6W9;A0A0B4LHY1;A8JRI1;Q7KRS7;Q8WR53;Q95UI6 | + | 2.915475539 | 0.003177305 | 1.060410261 |
| Flo2 | O61492;O61492-2;O61492-3 | + | 2.508104553 | 0.00618123 | 1.056554794 |
| Dmel\CG7322 | Q9VWP2 | + | 2.376314689 | 0.008758916 | 1.034769058 |
| RhoGAP19D | E1JJS2;M9NFH4;Q9VRA6;X2JEI9 | + | 2.898081805 | 0.003229698 | 1.03435564 |
| Ttd14 | Q7K556;Q7K556-2 | + | 2.540943614 | 0.005916248 | 1.034058094 |
| Dmel\CG16989 | Q9W5D3 | + | 4.589354845 | 0 | 1.034016132 |
| Nbr | Q9VIF1 | + | 2.297805263 | 0.009981208 | 1.032131195 |
| DmRH5 | Q8MZI3 | + | 2.937705358 | 0.00315122 | 1.028904915 |
| AMPdeam | M9MS55;Q961Q7;Q9VY76;X2JEY5 | + | 3.069309745 | 0.002816092 | 1.028597355 |
| galectin | E1JHP9;E1JHQ0;M9NCX9;Q9VPI6 | + | 3.414623389 | 0.001310044 | 1.02710104 |
| Dlic | Q9VZ20 | + | 2.462188079 | 0.007239567 | 1.023853302 |
| Root | Q9VCD1 | + | 3.067415589 | 0.002853868 | 1.019757509 |
| pyd | A0A0B4K6Y7;A8JQV8;Q9VHK1 | + | 2.382657033 | 0.008682927 | 1.017354012 |
| Wee1 | P54350 | + | 2.989843286 | 0.002903394 | 1.017009735 |
| nudC | Q9VVA6 | + | 2.790199647 | 0.003966667 | 1.015706539 |
| CG12096 | Q9VYG1 | + | 5.181646363 | 0 | 1.010892391 |
| RanBPM | A0A0B4KFL0;A0A126GUM4;Q4Z8K6 | + | 5.034740815 | 0 | 1.009589195 |
| Prosbeta7 | Q9VNA5 | + | 2.333526356 | 0.009495845 | 1.007863045 |
| Tbcc | Q95TP0 | + | 2.560101516 | 0.005837288 | 1.004814625 |
| Ate1 | O96539;O96539-2;O96539-4;O96539-5 | + | 3.675408895 | 0.001177778 | 0.997638464 |
| Dmel\CG10376 | Q9VJ61 | + | 2.427263017 | 0.007737313 | 0.996046782 |
| CG2200 | Q9VYH3 | + | 2.44619601 | 0.007495441 | 0.995422125 |
| trc | Q9NBK5-2 | + | 2.514692956 | 0.006166395 | 0.993583918 |
| Dmel\CG10465 | Q7JZ62 | + | 2.341069535 | 0.009454039 | 0.9896276 |
| Ire1 | A8JR46 | + | 2.840714496 | 0.003660131 | 0.981097937 |
| dos | Q9VZZ9;Q9VZZ9-2 | + | 2.63951202 | 0.005001815 | 0.979777813 |
| Dmel\CG9175 | Q9VMH4 | + | 4.207677965 | 0.00019802 | 0.975657463 |
| GMF | Q9VJL6 | + | 4.6031215 | 0 | 0.972860336 |
| tws | P36872;P36872-2 | + | 2.440142885 | 0.007625943 | 0.971441746 |
| mal | Q9VRA2 | + | 2.331846861 | 0.009456552 | 0.962864399 |
| P5cr | Q9V3F8 | + | 3.633658156 | 0.00112766 | 0.952661276 |
| p47 | Q7K3Z3 | + | 3.01477873 | 0.002784946 | 0.950202227 |
| Gmppb | Q7JZB4 | + | 2.470483384 | 0.00724493 | 0.949764252 |
| Prosbeta6 | P40304 | + | 3.233491305 | 0.002006969 | 0.949352026 |
| mrj | Q7JUZ6 | + | 2.459956846 | 0.007241911 | 0.947811604 |
| rngo | Q9VXF9 | + | 3.615457146 | 0.001104167 | 0.943232536 |
| GstD4;GstD7 | Q9VG93;Q9VG96 | + | 2.340494354 | 0.00944089 | 0.937876225 |
| Vha14-1 | Q24583 | + | 3.312053153 | 0.001463035 | 0.936879635 |
| Cdc37 | Q24276 | + | 4.567130592 | 0 | 0.9240098 |
| Ube3a | Q9VTH1 | + | 3.270014141 | 0.001665428 | 0.916270733 |
| CG17528 | Q7PLI7 | + | 3.187110276 | 0.001986928 | 0.911474466 |
| Mitf | C3KGP2 | + | 2.326380744 | 0.00943956 | 0.907089472 |
| CIAPIN1 | Q9V3Y2 | + | 3.321245622 | 0.001412698 | 0.904081821 |
| Drice | O01382 | + | 2.909361062 | 0.0032 | 0.899027348 |
| RagC-D | Q7K519 | + | 2.633768064 | 0.005059459 | 0.880333185 |
| Dmel\CG7326 | Q9VWP0 | + | 2.420174093 | 0.007952522 | 0.877381802 |
| Spn55B | Q7JV69 | + | 2.452183869 | 0.007389313 | 0.869242668 |
| Ubr3 | Q9W3M3;Q9W3M3-2 | + | 3.061709858 | 0.002829545 | 0.864006758 |
| Pde1c | B7YZV4;B7YZV4-2 | + | 2.79344639 | 0.003899791 | 0.862593412 |
| AP-2sigma | Q9VDC3 | + | 4.300607823 | 0 | 0.861504555 |
| Dmel\CG7600 | Q9XZ12 | + | 2.40546609 | 0.008233577 | 0.860736847 |
| Vps60 | Q9VVI9 | + | 2.810745814 | 0.003897655 | 0.849918127 |
| Lamtor5 | O96824 | + | 2.617077284 | 0.005252236 | 0.838781834 |
| UFSP1 | Q4V6M7 | + | 3.397454782 | 0.001378723 | 0.837348223 |
| Smox | O96660 | + | 2.928285753 | 0.003230769 | 0.826051235 |
| Tbcb | A1ZBM2 | + | 3.455011636 | 0.001285068 | 0.822036505 |
| Dmel\CG10973 | Q95RI2 | + | 2.860628813 | 0.003497758 | 0.818817616 |
| Plap | Q9VPY2 | + | 4.454883772 | 0 | 0.805756569 |
| Rpt5 | Q9V3V6 | + | 2.297416441 | 0.009967828 | 0.80343914 |
| RagA-B | Q9VHJ4 | + | 3.9522376 | 0.000603175 | 0.798233032 |
| Smyd4-4 | Q7KMH5 | + | 2.526720917 | 0.005966997 | 0.79306221 |
| LTV1 | Q7KN79 | + | 2.6548548 | 0.004985185 | 0.781851053 |
| Nubp1 | Q9VJI9 | + | 2.931237991 | 0.003174757 | 0.778158188 |
| Akt | Q8INB9 | + | 2.314033463 | 0.009686221 | 0.770503998 |
| mats | Q95RA8 | + | 2.985753783 | 0.002873385 | 0.73824358 |
| Ubr1 | Q9VX91 | + | 2.360579307 | 0.009100141 | 0.73631382 |
| Mettl2 | Q86BS6-3 | + | 2.504014932 | 0.006177134 | 0.724598646 |
| Tao | Q0KHQ5 | + | 2.305206667 | 0.009831522 | 0.720607281 |
| Dmel\CG10222 | Q9VU67 | + | 2.589632397 | 0.005425606 | 0.714279413 |
| MESK2 | Q8T0V2;Q9I7V6;Q9I7V7 | + | 2.530985861 | 0.005956882 | 0.699973345 |
| Naa35 | Q9W1A2;Q9W1A2-2 | + | 2.334800696 | 0.009464632 | 0.678515196 |
| anon-EST:Posey268 | A0A0B4LFP2;Q9W5T4 | + | 2.682977253 | 0.004613636 | 0.6730268 |
| Fkbp12 | P48375 | + | 2.994232961 | 0.002899471 | 0.650327206 |
| Cdk5 | P48609 | + | 3.186997119 | 0.001980456 | 0.641967773 |
| Oda | P54361 | + | 2.490174706 | 0.006647799 | 0.632522106 |
| Ranbp21 | Q9VWE7 | + | 3.381604967 | 0.001333333 | 0.618779659 |
| Ubc4 | P52486 | + | 2.534837071 | 0.00596 | 0.612205267 |
| Fkbp59 | Q9VL78 | + | 2.560897022 | 0.005847199 | 0.610409498 |
| DmCG31729 | X2JDZ6 | + | 2.302987136 | 0.009989189 | 0.5978055 |
| Dmel\CG1943 | Q9VI56 | + | 2.743295226 | 0.00410297 | 0.588564873 |
| lic | O62602 | + | 3.098312231 | 0.002592375 | 0.567937374 |
| Elp5 | Q24050 | + | 2.605309883 | 0.005206349 | 0.56756258 |
| alph | Q8IMK7;Q961C5;Q9VAK1 | + | 2.970452371 | 0.002997468 | 0.564542055 |
| Rox8 | Q8IMX4;Q9VCE3 | + | 2.426713295 | 0.007738095 | 0.55867815 |
| Trx-2 | Q9V429 | + | 2.800394027 | 0.003890295 | 0.554726124 |
| cin | P39205 | + | 2.414969578 | 0.008105882 | 0.551912308 |
| fus | Q9BJZ5 | + | 2.767757178 | 0.004065306 | 0.547292233 |
| DnaJ-1 | Q24133 | + | 2.731154704 | 0.004156556 | 0.535257339 |
| EDTP | A1ZAS8;Q9NDR1 | + | 3.957332927 | 0.000608 | 0.5042274 |
| Npl4 | Q9VBP9;Q9VBP9-2 | + | 2.387885488 | 0.008571429 | 0.488253593 |
| PDCD-5 | Q9VUZ8 | + | 2.911999285 | 0.003169811 | 0.487480402 |
| Dmel\CG12773;EG:8D8.3 | M9PDG9;O46100 | + | 2.498220828 | 0.006351438 | 0.301655054 |
