## Supplementary Table 2 for "Vesicular pseudopodia define the fusion site on large secretory vesicles of the *Drosophila* salivary glands"

| **Line name** | **Genotype (for commercial lines)** | **Source/accession number** |
| --- | --- | --- |
| UAS-LifeAct-Ruby | y[1] w[*]; P{y[+t*] w[+mC]=UAS-Lifeact-Ruby}VIE-19A | BDSC_35545 |
| Sgs3-GFP | w[*]; P{w[+mC]=Sgs3-GFP}2 | BDSC_5884 |
| Sgs3-DsRed | P{w^+^, Sgs3-DsRed} | Kind gift from Julie Brill |
| C135-GAL4 | w[1118]; P{w[+mW.hs]=GawB}path[c135] | BDSC_6978 |
| fkh-GAL4 | w[*]; P{w[+mC]=fkh-GAL4.H}3 | BDSC_78060 |
| UAS-MIM-Emerald |  | Biton et al 2024 |
| UAS-MIM-mScarlet |  | Biton et al 2024 |
| UAS-IVS-myrtdTomato | w[*]; P{y[+t7.7] w[+mC]=10XUAS-IVS-myr::tdTomato}attP2 | BDSC_32221 |
| UAS-GFP-Sec15 | w[*]; P{w[+mC]=UAS-GFP.Sec15}2/CyO | BDSC_39685 |
| Sec15-sfGFP-TVPTBF | FlyFos015736(pRedFlp-Hgr)(sec15[32881]::2XTY1-SGFP-V5-preTEV-BLRP-3XFLAG)dFRT | VDRC_318013 |
| UASp-CD63-GFP | w[*]; P{w[+mC]=UAS-EGFP.CD63}2; Dr[1]/TM3, Sb[1] | Kind gift from Eli Arama |
| UASt-CD63-GFP | w[*]; P{w[+mC]=UAS-EGFP.CD63}2; Dr[1]/TM3, Sb[1] | BDSC_91390 |
| Sqh-mCherry | w[*]; P{w[+mC]=sqh-mCherry.M}3 | BDSC_59024 |
| Tsp42Ee-EGFP-polyhis | y[1] w[*]; P{w[+mC]=PTT-GC}Tsp42Ee[CC01420] | BDSC_51558 |
| UAS-Tsp42Ef-3xHA | M{UAS-Tsp42Ef.ORF.3xHA.GW}ZH-86Fb | F004035 |
| UAS-Tsp29Fa |  | Kind gift from Julie Brill |
| UAS-Tsp96F-3xHA | M{UAS-Tsp96F.ORF.3xHA.GW}ZH-86Fb | F003566 |
| Df(2R)BSC260 | w[1118]; Df(2R)BSC260/CyO | BDSC_23160 |
| MIMnull |  | Kind gift from Helen Zenner |
| Tsp42Ee KD#1 | y[1] v[1]; P{y[+t7.7] v[+t1.8]=TRiP.HMJ23660}attP40/CyO | BDSC_62303 |
| Tsp42Ee KD#2 | y[1] sc[*] v[1] sev[21]; P{y[+t7.7] v[+t1.8]=TRiP.HMC06505}attP40 | BDSC_77368 |
